# A Generalized Framework for the Calcium Control Hypothesis Describes Weight-Dependent Synaptic Plasticity

**DOI:** 10.1101/2023.07.13.548837

**Authors:** Toviah Moldwin, Li Shay Azran, Idan Segev

## Abstract

The brain modifies synaptic strengths to store new information via long-term potentiation (LTP) and long-term depression (LTD). Evidence has mounted that long-term plasticity is controlled via concentrations of calcium ([Ca^2+^]) in postsynaptic spines. Several mathematical models describe this phenomenon, including those of Shouval, Bear, and Cooper (SBC) (Shouval et al., 2002, 2010) and Graupner and Brunel (GB)(Graupner & Brunel, 2012). Here we suggest a generalized version of the SBC and GB models, based on a fixed point – learning rate (FPLR) framework, where the synaptic [Ca^2+^] specifies a fixed point toward which the synaptic weight approaches asymptotically at a [Ca^2+^]-dependent rate. The FPLR framework offers a straightforward phenomenological interpretation of calcium-based plasticity: *the calcium concentration tells the synaptic weight where it is going and how fast it goes there*. The FPLR framework can flexibly incorporate various experimental findings, including the existence of multiple regions of [Ca^2+^] where no plasticity occurs, or plasticity in cerebellar Purkinje cells, where the directionality of calcium-based synaptic changes is thought to be reversed relative to cortical and hippocampal neurons. We also suggest a modeling approach that captures the dependency of late-phase plasticity stabilization on protein synthesis. We demonstrate that due to the asymptotic, saturating nature of synaptic changes in the FPLR rule, the result of frequency- and spike-timing-dependent plasticity protocols are weight-dependent. Finally, we show how the FPLR framework can explain plateau potential-induced place field formation in hippocampal CA1 neurons, also known as behavioral time scale plasticity (BTSP).

## Introduction

Since the work of Donald Hebb (Hebb, 1949), it has been believed that the brain learns via modifying the strengths of synaptic connections between neurons. Decades of experimental research have shown that synaptic strengths can be increased via a process known as long-term potentiation (LTP) and decreased via another process known as long term depression (LTD). Experimentally, there are a variety of protocols that can induce either potentiation or depression, usually involving stimulating the presynaptic inputs (e.g., via electrical or optical stimulation of the axon, or glutamate uncaging at the synapse), the postsynaptic soma (e.g., by electrically inducing subthreshold depolarization or spiking), or both, for varying frequencies and durations (Shouval et al., 2010).

Over the past several decades, plasticity researchers have been drawn to the possibility that there may be a fundamental molecular process underlying long-term plasticity. Lisman (J. Lisman, 1989) proposed a calcium-based theory of plasticity, a theory which has subsequently been extensively validated (Cho et al., 2001; Cummings et al., 1996; J. Lisman, 1989; Shouval et al., 2002; Yang et al., 1999). In this framework, different presynaptic and postsynaptic stimulation protocols yield potentiation or depression due to the calcium concentration ([Ca^2+^]) which is induced at the synapse by the stimulation. Specifically, if the [Ca^2+^] is low, no change occurs. If the [Ca^2+^] rises above a critical threshold for depression (*θ*_*D*_), long-term depression (LTD) occurs and the synaptic strength is decreased. If the [Ca^2+^] is above the critical threshold for potentiation (*θ*_*P*_), long-term potentiation (LTP) occurs and the synaptic strength is increased. It is believed that calcium promotes LTP via pathways involving protein kinases such as calmodulin kinase (CaMKII) (J. Lisman, 1989; Malenka et al., 1989; Neveu & Zucker, 1996; R et al., 1989), while promoting LTD via phosphatases such as calcineurin (J. Lisman, 1989; Mulkey et al., 1994; RM et al., 1993).

The success of the calcium theory of plasticity has attracted theoretical neuroscientists and modelers to attempt to capture the dynamics of calcium-based plasticity in equations and computer models. There are two popular models of calcium-based plasticity that are currently in use: the models of Shouval, Bear, and Cooper (SBC)(Shouval et al., 2002, 2010) and Graupner and Brunel (GB)(Graupner & Brunel, 2012). There are several differences between these models, and we will explore them at length. Fundamentally, though, both models attempt to capture 3 essential properties:

1. Synaptic weights decrease when the synaptic [Ca^2+^] is in the depressive calcium region (*θ*_*D*_ ≤ [*Ca*^2+^] ≤ *θ*_*P*_) and potentiate when the [Ca^2+^] is in the potentiative calcium range ([*Ca*^2+^] > *θ*_*P*_).
2. Synaptic weights do not potentiate or decrease linearly *ad infinitum* in the presence of potentiating or depressing levels of calcium, but rather they increase or decrease asymptotically toward some maximum or minimum value. This accounts for the fact that in biology synaptic weights cannot be arbitrarily large or arbitrarily small (excitatory synapses can’t have negative weights) and the fact that synaptic strengths are observed to exhibit saturating behavior when undergoing a plasticity protocol (O’Connor et al., 2005b).
3. In the pre-depressive region of [Ca^2+^], neither potentiation or depression occur, but synaptic strengths may still “drift” toward some pre-determined value(s), called stable states or fixed points, over a long time scale (i.e. hours or days). This feature is motivated by evidence that synapses may have a discrete number of stable states and that synaptic strengths have been experimentally observed to slowly drift toward stable states after inducing potentiation or depression strengths (Frey & Morris, 1997; Kauderer & Kandel, 2000; Manahan-Vaughan et al., 2000; Redondo et al., 2010; Sajikumar et al., 2005). The major practical difference between the SBC model and the GB model pertains to the drift in the pre-depressive region of calcium. In the SBC model, weights drift towards a single baseline whereas in the GB model, synapses that are above a certain weight are stabilized toward an UP state while synapses that are below that weight are stabilized toward a DOWN state.

In this work, we will review and compare the SBC and GB models and make several modifications to both models to make them more analytically straightforward, thus allowing both experimentalists and theoreticians to use them with greater ease. Our modifications are based on the principle that a calcium-based plasticity model should allow for explicit and independent specification of critical parameters that capture the essential dynamics of the plasticity. Specifically, it should be possible to specify the fixed points of the asymptotic dynamics of the synaptic weights in each region of the [Ca^2+^] space (and in each region of the weight space in the GB rule), and it should also be possible to independently specify the rates (time constants) at which potentiation, depression, and drift take place without constraining the fixed points. We therefore call our model the fixed point – learning rate (FPLR) framework, as its dynamics can be fully characterized by specifying the fixed points and learning rates (time constants).

The FPLR framework is a phenomenological—as opposed to mechanistic—model of plasticity. The underlying assumption is that different levels of calcium specify different processes in the cell that have different teleological endpoints (i.e., the fixed points of the weights) and that occur at different rates. In other words, *the calcium concentration tells the synaptic weight where it is going and how fast it goes there*.

The dynamics of calcium-based plasticity can be captured by performing experiments where the synaptic [Ca^2+^] is held at a fixed level for some duration of time (on the order of minutes) until the synaptic weight reaches an asymptote, and then decreased to 0 for a longer duration of time (on the order of hours), which reveals both the fixed points and the rate at which they are reached (Fig. 1A).

**Figure 1:**
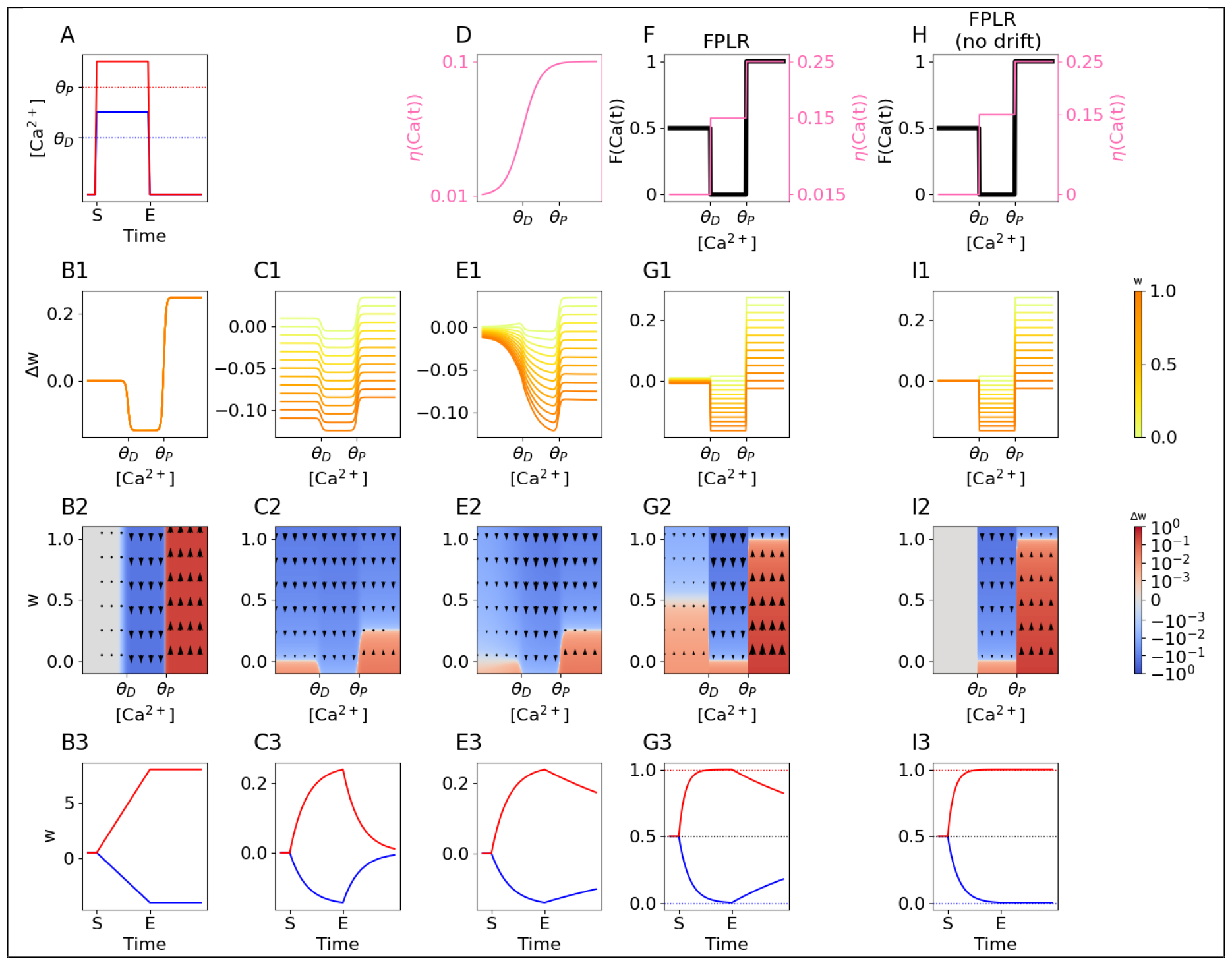
SBC and FPLR Rules for Calcium-based plasticity. **(A)** Basic stimulation to test the plastic effect of different levels of [*Ca*^2+^]. [*Ca*^2+^] is either raised to a depressive level (*θ*_*D*_ ≤ [*Ca*^2+^] ≤ *θ*_*P*_, blue line) or to a potentiative level ([*Ca*^2+^] > *θ*_*P*_, red line) for several minutes and is then reduced to and held at 0 for several hours to observe calcium-independent drift effects. **(B1)** Basic SBC two-threshold rule for [*Ca*^2+^]-dependent weight changes. **(B2)** Same as (B1) but with Δ*w* presented as a heatmap of both the present value of *w* and the [*Ca*^2+^]. In the basic SBC rule, there is no dependence on the present value of *w* and thus no variation along the vertical axis. Colors are log-normalized for visualization purposes. Red indicates potentiation, blue indicates depression, white indicates no change. **(B3)** Resultant weights over time when applying the basic SBC rule to the stimulation protocol from (A). Red and blue lines correspond to the red and blue lines (potentiative and depressive protocols, respectively) from (A). In the basic SBC rule, synapses potentiate or depress linearly in the presence of calcium and remain stable in the absence of calcium. **(C1)** SBC rule with weight decay. Here Δ*w* depends on both the present value of *w* and the [*Ca*^2+^]. Larger weights (darker lines) depress faster and potentiate more slowly than lower weights (lighter lines, some weight values are negative for illustrative purposes). **(C2)** Same as (C1), presented as a heatmap. White line indicates fixed points. **(C3)** Weights over time in the SBC rule with weight decay. Solid lines and dashed lines indicate different weight initializations. Dotted horizontal lines indicate fixed points. **(D)** Calcium-dependent learning rate in the original SBC rule. The rate of weight change increases sigmoidally with the [*Ca*^2+^], allowing for a slower drift to baseline for pre-depressive levels of [*Ca*^2+^]. **(E1-E3)** Dynamics of SBC rule with weight decay and [*Ca*^2+^]-dependent learning rate. **(F)** Fixed points (*F*(*Ca*(*t*)), black line, left y-axis) and learning rates (*η*(*Ca*(*t*)), pink line, right y-axis) for each region of [*Ca*^2+^] as step functions in the FPLR rule. **(G1-G3)** Dynamics of the FPLR rule. **(H)** Fixed points and learning rates for the modified SBC rule without drift. *η*(*Ca*(*t*)) is set to 0 in the pre-depressive region of [*Ca*^2+^]. **(I1-I3)** Dynamics of the FPLR rule with no drift.

The FPLR framework is more generic than the original SBC and GB rules, making it easy to specify arbitrary fixed points and learning rates based on experimental results. We show how the FPLR framework can be used to explore the dynamics of experimental results that are not captured by the original SBC and GB rules, such as incorporating two additional regions of [Ca^2+^] where neither potentiating nor depressive mechanisms are active, based on (Cho et al., 2001; J. E. Lisman, 2001; Tigaret et al., 2016), and modeling plasticity in Purkinje cells where the order of the potentiative region and depressive region are switched (i.e. *θ*_*D*_>*θ*_*P*_), based on (Coesmans et al., 2004; Piochon et al., 2016).

To synthesize the SBC and GB rules in the FPLR framework, we take a first step at expanding [Ca^2+^]-based plasticity models to encompass protein synthesis-dependent late-LTP (L-LTP) by proposing a framework where the [Ca^2+^]-dependent rules are conditional on the presence of synaptic proteins, and suggest that plasticity may behave like the SBC rule (drift to baseline) in the absence of protein synthesis, and more like the GB rule (stabilization) when protein synthesis has occurred.

An important aspect of the FPLR framework, as well as the original GB and SBC rules, is that because the dynamics of plastic changes are asymptotic, the magnitude (and sometimes direction) of plastic change in response to a given plasticity protocol will depend on the initial synaptic weight. We illustrate this by reproducing simple frequency dependent plasticity and spike-timing-dependent-plasticity (STDP) protocols and demonstrating how different initial weights lead to different plastic consequences in the FPLR framework.

Finally, to demonstrate a novel application of the FPLR rules, we propose a calcium-based solution to the mechanism underlying plateau potential-induced place field formation in the hippocampus, also known as behavioral time scale plasticity (BTSP) (Bittner et al., 2015, 2017; Milstein et al., 2021). We show that the FPLR framework can explain how a plateau potential can either “overwrite” a previously created place field or create a new place field without overwriting the first place field, depending on the distance between the new and old place field.

## Results

### Basic SBC Rule

We first consider the calcium-based plasticity rules of Shouval et al. (Shouval et al., 2002), starting with the simplest formulation. In this rule, synaptic strength is modified in a straightforward manner according to the depression and potentiation thresholds: at any time step *t*, if the calcium concentration ([Ca^2+^]) is in the pre-depressive range ([*Ca*^2+^] < *θ*_*D*_) the synaptic weight *w* remains unchanged. If the calcium concentration is in the depressive range (*θ*_*D*_ ≤ [*Ca*^2+^] ≤ *θ*_*P*_) *w* is decreased, and if the [Ca^2+^] is in the potentiating range ([*Ca*^2+^] > *θ*_*P*_), *w* is increased. Formally, the change in the synaptic weight Δ*w* (Δ*w* = *w*(*t* + 1) − *w*(*t*)) at a given time as a function of the local [Ca^2+^] at that time (denoted as *Ca*(*t*)) is given by:

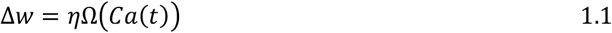

Where *η* is the learning rate and Ω is the two-threshold calcium based plasticity function described above, which can be expressed most simply as a step function:

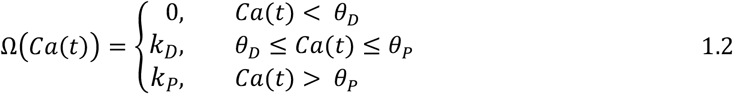

Where *k*_*D*_and *k*_*P*_ are the signed rates of depression and potentiation, respectively (*k*_*D*_< 0, *k*_*P*_> 0), and *θ*_*D*_and *θ*_*P*_represent the thresholds for depression and potentiation. If smooth transitions between regions are desired, this can also be expressed with a soft threshold using the sum of sigmoids (slightly modified from (Shouval et al., 2002)):

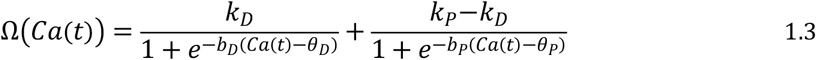

where *b*_D_and *b*_P_control the sharpness of the transitions between regions in the Ω function.

In the basic SBC rule, synaptic weights increase linearly at a rate of *k*_*P*_ in the potentiative region of [*Ca*^2+^], decrease linearly at rate of *k*_*D*_in the depressive region of and remain stable in the pre-depressive region of (Fig. 1B).

(We note that it may be more biologically plausible to implement the plasticity as a delayed function of the calcium signal, in which case one may substitute *Ca*(*t* − *D*) in place of *Ca*(*t*), where *D* indicates the duration of the temporal delay between the calcium signal and the plastic effect. For simplicity, however, we will assume that there is no such delay, i.e. *D* = 0.)

### SBC Rule with Weight Decay

To prevent synapses becoming arbitrarily large or small, Shouval et al.(Shouval et al., 2002) added a weight decay term to the original plasticity rule.

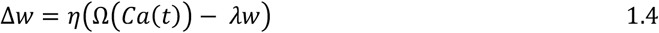

Where *λ* is the rate of decay. Importantly, the change in the weight Δ*w* now depends on both the [Ca^2+^] and the present value of the weight *w*. It can be instructive to visualize Δ*w* as a function of both *w* and [Ca^2+^]. We show (Fig. 1C) how the SBC rule with weight decay differs from the rule without it. In the pre-depressive range of [Ca^2+^], *w* decreases (or increases, if it is negative) asymptotically to the fixed point of 0 at a rate of *η*λ . In the depressive [Ca^2+^] range, *w* decreases asymptotically to the fixed point of 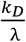 (or increases if the weight is below that point). In the potentiating range of [Ca^2+^], *w* increases asymptotically to the fixed point of 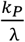 (or decreases if the weight is above the fixed point). Note that in this framework, there is no way to independently change the asymptotic behavior without changing either the rates of potentiation and depression (*k*_*D*_ and *k*_*P*_) or the decay rate λ.

One way to visualize the dynamics of the SBC plasticity rule with and without weight decay is the simple experiment mentioned above where the synapse is exposed to a fixed amount of calcium in either the depressive or potentiating range over some time period, after which the calcium is decreased to 0 (Fig. 1A). In the original SBC rule without weight decay, the weight linearly increases (or decreases) for as long as the calcium pulse is present, then immediately stops changing (i.e., remains stable) when the calcium is turned off (Fig. 1B3). By contrast, when weight decay is used, the synapse potentiates or depresses asymptotically toward the fixed points when the calcium pulse is on, then drifts toward 0 when the calcium is turned off (Fig. 1C3).

### SBC Rule with Weight Decay and Ca^2+^-dependent learning rate

Weight decay that drifts quickly toward 0 in the absence of plasticity-inducing calcium may be an undesirable feature of a plasticity rule if we wish to model synapses that are potentiated or depressed for a long duration. Shouval et al. (Shouval et al., 2002) therefore introduced a sigmoidal [Ca^2+^]-dependent learning rate, Ω(*Ca*(*t*)), to mitigate the effect of the weight decay in the absence of calcium. The SBC rule with the calcium-dependent learning rate is thus defined as:

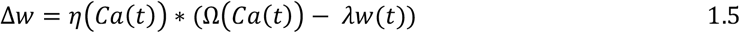

The basic idea is that instead of having a constant learning rate, the learning rate (including the rate of the weight decay) increases in a sigmoidal fashion with the amount of calcium (Fig. 1D). Thus, at pre-depressive levels of [Ca^2+^] the weight will decay slowly, allowing for greater stability over long time horizons. (Fig. 1E).

### Fixed point – learning rate version of the SBC Rule

In the FPLR framework, we propose a modified version of the SBC rule which allows the modeler to specify the fixed points and learning rates explicitly in all three region of [Ca^2+^]:

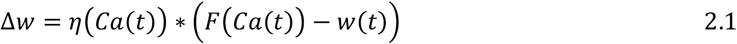

Here, instead of Ω(*Ca*(*t*)), we use *F*(*Ca*(*t*)), a step function which describes the *fixed points* of the weights as a function of the [Ca^2+^]. (We note that if *λ* is fixed to 1 in the original SBC rule [equation 1.5], Ω(*Ca*(*t*)) also specifies the fixed points of the weights, however *F*(*Ca*(*t*)) is defined this way explicitly.) Moreover, *η*(*Ca*(*t*)) here is also a 3-valued step function (as opposed to a sigmoid in the original SBC rule) which determines the rate at which the weight asymptotically approaches the fixed point for each calcium level. (We require *η*(*Ca*) ≤ 1 for all values of *Ca* to prevent oscillations). In this framework, the learning rate *η*(*Ca*(*t*)) defines the fraction of the distance between the current weight and the fixed point *F*(*Ca*(*t*)) which is traversed at each time step. (If *η*(*Ca*(*t*)) = 1) the synapse immediately jumps to the fixed point specified by *F*(*Ca*(*t*)), which can be useful for modeling discrete-state synapses.)

For example, we might have:

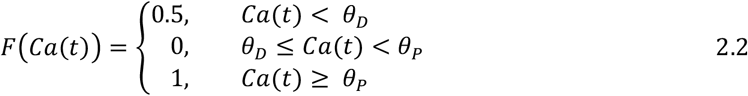

And

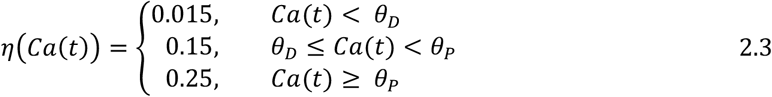

(Fig 1F) This means that synapses with pre-depressive calcium concentrations eventually drift toward a “neutral” state of 0.5 at a rate of 0.015, synapses with a depressive calcium concentration will depress towards 0 at a rate of 0.15, and synapses with a potentiative [Ca^2+^] will be potentiated towards 1 at a rate of 0.25 (Fig. 1G). (See (Enoki et al., 2009) for experimental evidence that synapses at baseline can be either potentiated or depressed). (Note that the fixed points and rates used here and in subsequent sections are specified in arbitrary units and meant to illustrate qualitative dynamics of the synapse only; see Figure 5 and **Methods** for biologically plausible parameters).

We can also turn off the drift in the pre-depressive region entirely by setting *η*(*Ca*(*t*) < *θ*_*D*_) = 0, thus allowing for synapses that are stable at every weight unless modified by depressive or potentiative calcium concentrations (Fig 1H,1I). This version of the rule can be useful for contexts in which the modeler is primarily interested in understanding the early phase of plasticity, or for theoretical work exploring the consequences of synaptic weights that potentiate and depress asymptotically.

### Versatility of the FPLR framework

Until now, all of the above rules have assumed that there are only three regions of [Ca^2+^] relevant for plasticity and that *θ*_*D*_ < *θ*_*P*_. There are some experimental results that complicate this picture. For example, there is evidence that in Purkinje neurons, *θ*_*D*_ > *θ*_*P*_, and that it is possible to shift the plasticity thresholds via inputs from other neurons (Coesmans et al., 2004). This can also easily be implemented in our modified plasticity rule by changing the fixed points in each [Ca^2+^] region. (Fig. 2A,B)

**Figure 2:**
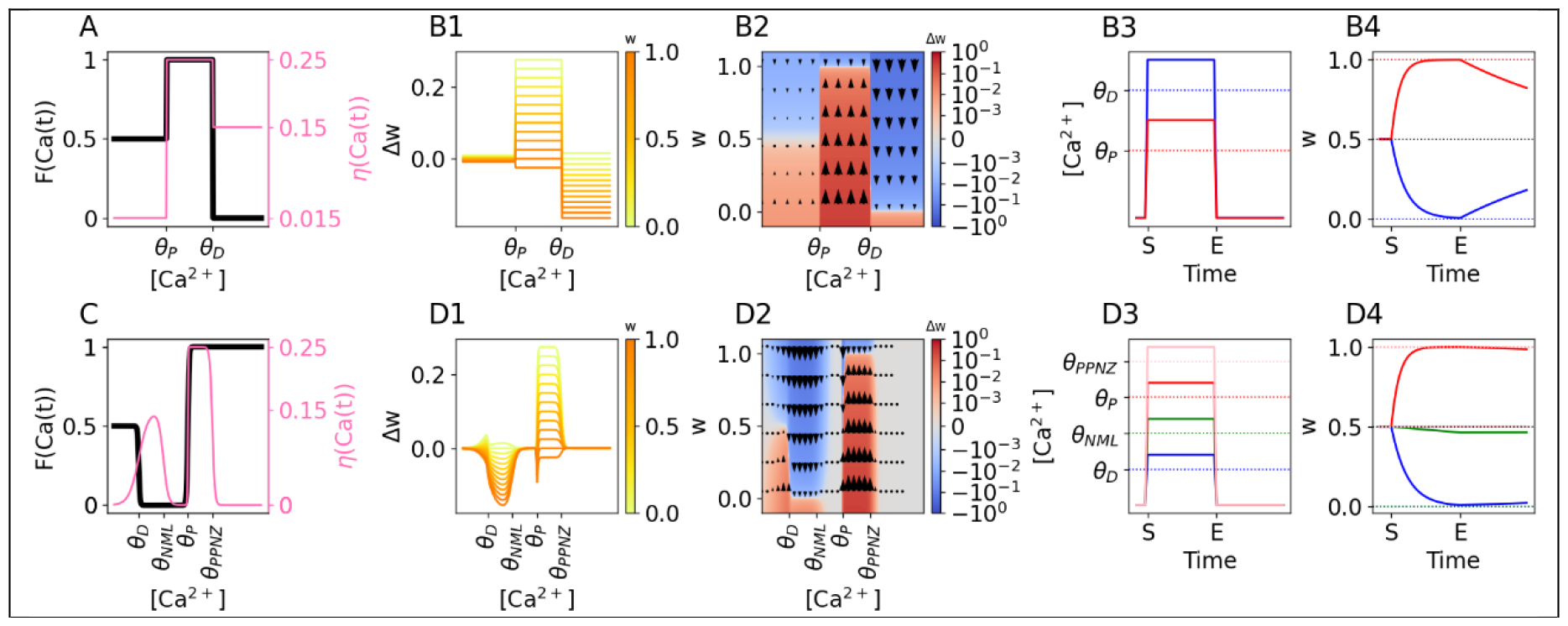
Incorporating Multiple Thresholds with the FPLR Rule. **(A)** Possible fixed points and learning rates for Purkinje cells, where the potentiation threshold is lower than the depression threshold (*θ*_*P*_< *θ*_*D*_).**(B1)** Δ*w* in modified SBC rule for Purkinje cells using the fixed points and learning rates from (A). **(B2)** Same as (B1), presented as a heatmap. **(B3)** Stimulation protocol for Purkinje cells. Here, the [*Ca*^2+^] used for potentiation (red line) is less than the [*Ca*^2+^] used to induce depression. **(B4)** Weights over time in Purkinje cells for the stimulation protocols in (B3). Dotted lines represent fixed points. Solid and dashed lines represent different weight initializations. **(C)** Fixed points and learning rates, defined using a soft-threshold step function, with two additional regions of [*Ca*^2+^] where *η* = 0 and therefore no plasticity is induced. *θ*_*NML*_: no man’s land, *θ*_*PPNZ*_: post-potentiative neutral zone. **(D1)** Δ*w* in modified SBC rule using the fixed points and learning rates from (C). Note the U-shaped dependence of depression magnitude on the [*Ca*^2+^] resulting in a no mans zone between the depressive and potentiative regions of [*Ca*^2+^]. Irregularities near the boundaries of [*Ca*^2+^] regions are due to the soft transitions in the fixed point and plasticity rate functions. **(D2)** Same as (D1), presented as a heatmap. **(D3)** Stimulation protocol for each region of [*Ca*^2+^] shown in (F). **(D4)** Weights over time for the stimulation protocols in (D3). Note that when the [*Ca*^2+^] is in the no man’s zone (green line) depression occurs at a much slower rate, and in the post-potentiative neutral zone (black line) nearly no plasticity occurs.

Moreover, even within hippocampal and cortical cells, there may be additional regions of [Ca^2+^] where the plasticity dynamics change. Cho et al. (Cho et al., 2001) found that within the depressive region of [Ca^2+^], the magnitude of depression exhibits a U-shaped relationship with the calcium concentration, such that a “No man’s zone” appears at the boundary between the depressive and potentiative region of [Ca^2+^], where the [Ca^2+^] is too large to induce depression but too small to induce potentiation (Cho et al., 2001; J. E. Lisman, 2001). It may make sense for a modeler to include this no-man’s zone more explicitly in the plasticity rule. There is also evidence from Tigaret et al. (Tigaret et al., 2016) that there is some maximum level of [Ca^2+^] beyond which potentiation mechanisms are inactivated. It would be worthwhile to be able to incorporate this “post-potentiative neutral zone” as well. These “neutral zones” can easily be incorporated into the modified SBC rule by adding additional thresholds into the step function from Eq. 2.1 and setting the learning rate to 0 in those regions (the fixed points in the no-plasticity regions can be chosen arbitrarily, as they are irrelevant if the learning rate in those regions is 0).

If desired, we can also explicitly model the U-shaped dependency between [Ca^2+^] and synaptic depression. Although we have used a hard threshold step function to implement the previous examples of the modified SBC rule, in principle both *F*(*Ca*(*t*)) and *η*(*Ca*(*t*)) can be arbitrary functions as long as *η*(*Ca*(*t*)) < 1 for all values of *Ca*(*t*). To implement a U-shaped region, we can extend the soft threshold step function equation from Eq. 1.3 into a general form to include arbitrary regions of [Ca^2+^]:

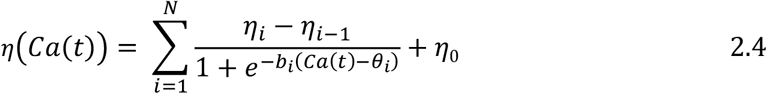

Where *θ*_1_ … *θ*_*i*_ … *θ*_*N*_ are the thresholds that differentiate between regions of [Ca2+] ordered such that *θ*_*i*_< *θ*_*i*+1_,*b*_1_… *b*_*i*_… *b*_*N*_ determine the steepness of transition between each [Ca2+] region, *η*_1_ … *η*_i_ … *η*_*N*_ are the learning rates in each [Ca2+] region, and *η*_0_ is the learning rate in the region between 0 and *θ*_1_. This representation approaches the equivalent step function as the values for *b*_*i*_become sufficiently large. (A similar equation can be used for *F*(*Ca*(*t*))). Using Eq. 2.4, we can easily create the U-shaped relationship between [Ca^2+^] and synaptic depression (Fig. 2C,D).

### Graupner and Brunel Model

One drawback of the SBC plasticity rules is that even in the final version with a calcium-dependent learning rate, synaptic weights eventually trend toward 0 in the presence of pre-depressive levels of calcium. There is some experimental evidence, however, that synapses are bistable, existing in a potentiated (UP) state with weight *w*_*UP*_ or a depressed (DOWN) state with weight *w*_*DOWN*_, and that synaptic strengths slowly trend toward one of those two states depending on the early synaptic strength after inducing a plasticity protocol (Bagal et al., 2005; O’Connor et al., 2005a; Petersen et al., 1998) . Graupner and Brunel (Graupner & Brunel, 2012) proposed a model that captured these dynamics. In their model, the synaptic efficacy, *ρ* (*ρ* is linearly mapped to *w* according to the equation *w* = *w*_*DOWN*_ + *ρ*(*w*_*UP*_ − *w*_*DOWN*_)) asymptotically decreases to the DOWN state (at *ρ* = 0) in the presence of depressive [Ca^2+^] or increases asymptotically to the UP state (near *ρ* = 1) in the presence of potentiating [Ca^2+^]. When the [Ca^2+^] is pre-depressive, *ρ* either increases toward 1 or decreases toward 0 in a hyperbolic fashion on a very slow time scale depending only on the present value of *ρ*. Specifically, if *ρ* is larger than the value of an unstable fixed point *ρ*_⋆_(set to 0.5), *ρ* trends toward the UP state whereas if *ρ* is smaller than this value, *ρ* trends toward the DOWN state (Fig. 3 - blue lines, Fig. 4A). Formally, we have:

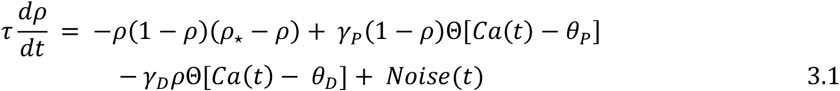

**Figure 3:**
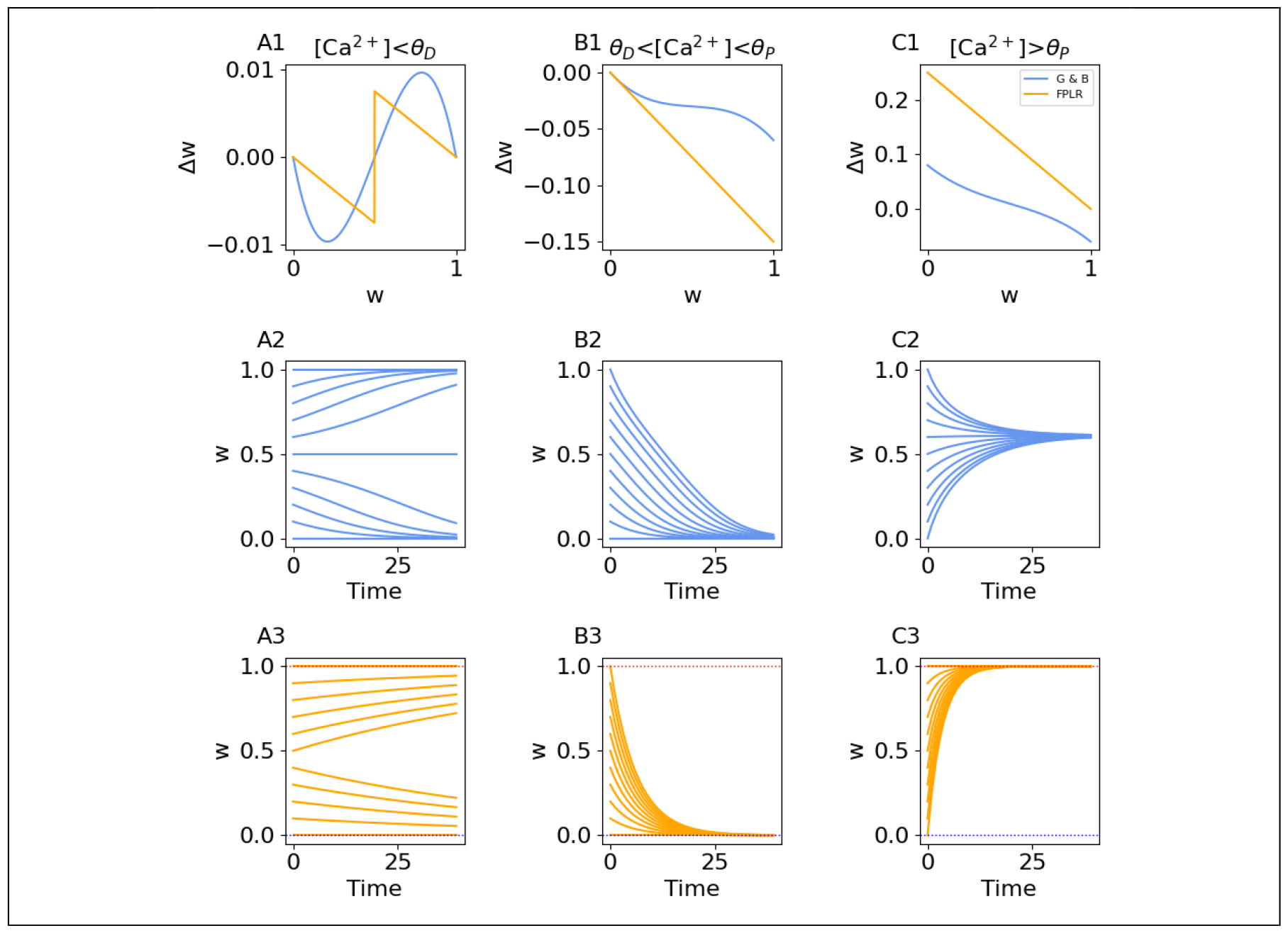
Simplifying the Original GB Rule. **(A1)** Δ*w* as a function of *w* for the original GB rule (solid blue line) and for the simplified GB rule (dashed orange line) in the pre-depressive region of [*Ca*^2+^]. **(A2)** Weights over time in the original GB rule in the pre-depressive region of [*Ca*^2+^] for synapses with different initial weights. Weights increase and decrease sigmoidally. **(A3)** Weights over time in the simplified GB rule, as in (A2). Weights increase and decrease asymptotically. **(B1-B3)** as in A for the depressive region of [*Ca*^2+^]. **(C1-C3)** As in A for potentiative region of [*Ca*^2+^].

**Figure 4:**
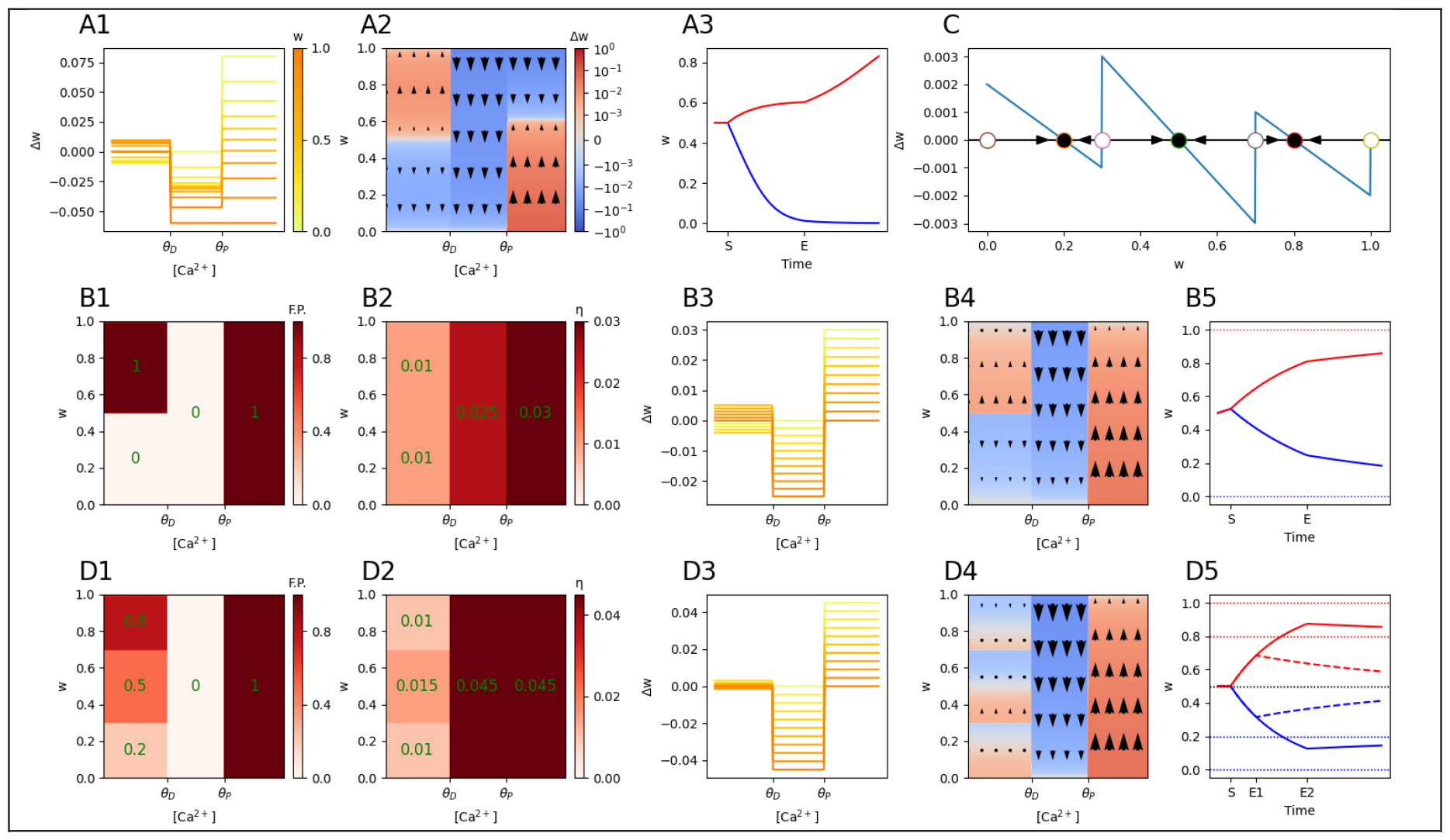
Comparing GB Rules. **(A1)** Δ*w* in the original GB rule (darker lines indicate larger values for the current weight). Note that in the pre-depressive region of [*Ca*^2+^], the weight may increase or decrease (i.e. Δw can be greater than or less than 0) depending on the value of the initial weight. **(A2)** Same as (A1), presented as a heatmap. **(A3)** Weights over time in the original GB rule for the stimulation protocols from Fig. 1A. Weights drift towards either the UP or DOWN stable states depending on their value when the [*Ca*^2+^] is reduced to 0. **(B1)** Fixed points for the two-dimensional FPLR rule as a function of the current weight and [*Ca*^2+^]. **(B2)** Learning rates for the two-dimensional FPLR rule **(B3-B4)** As in A1-A2 for the FPLR rule. **(B5)** Weights over time in the FPLR rule for the stimulation protocols from Fig. 1A. **(C)** Defining fixed points and basins of attraction in the generic GB rule. Within each region of [*Ca*^2+^], we can define an arbitrary number of stable fixed points (filled circles) as a function of the current weight *w*, as well as the boundaries of their respective non-overlapping basins of attraction (open circles). **(D1-D4)** As in (B) for an FPLR rule with a tristable synaptic weight in the pre-depressive drift region of [*Ca*^2+^], using fixed points from (C). **(D5)** A step of calcium of potentiative or depressive [*Ca*^2+^] is applied for either a short (2 seconds, dash-dot line, stimulation starts at S and ends at E1), or long (20 seconds, solid line, stimulation starts at S and ends at E2) duration. The long-duration stimulus escapes the middle fixed point’s basin of attraction; the short duration stimulus does not. Note that the long-duration stimulus overshoots/undershoots the late-phase fixed points.

**Figure 5:**
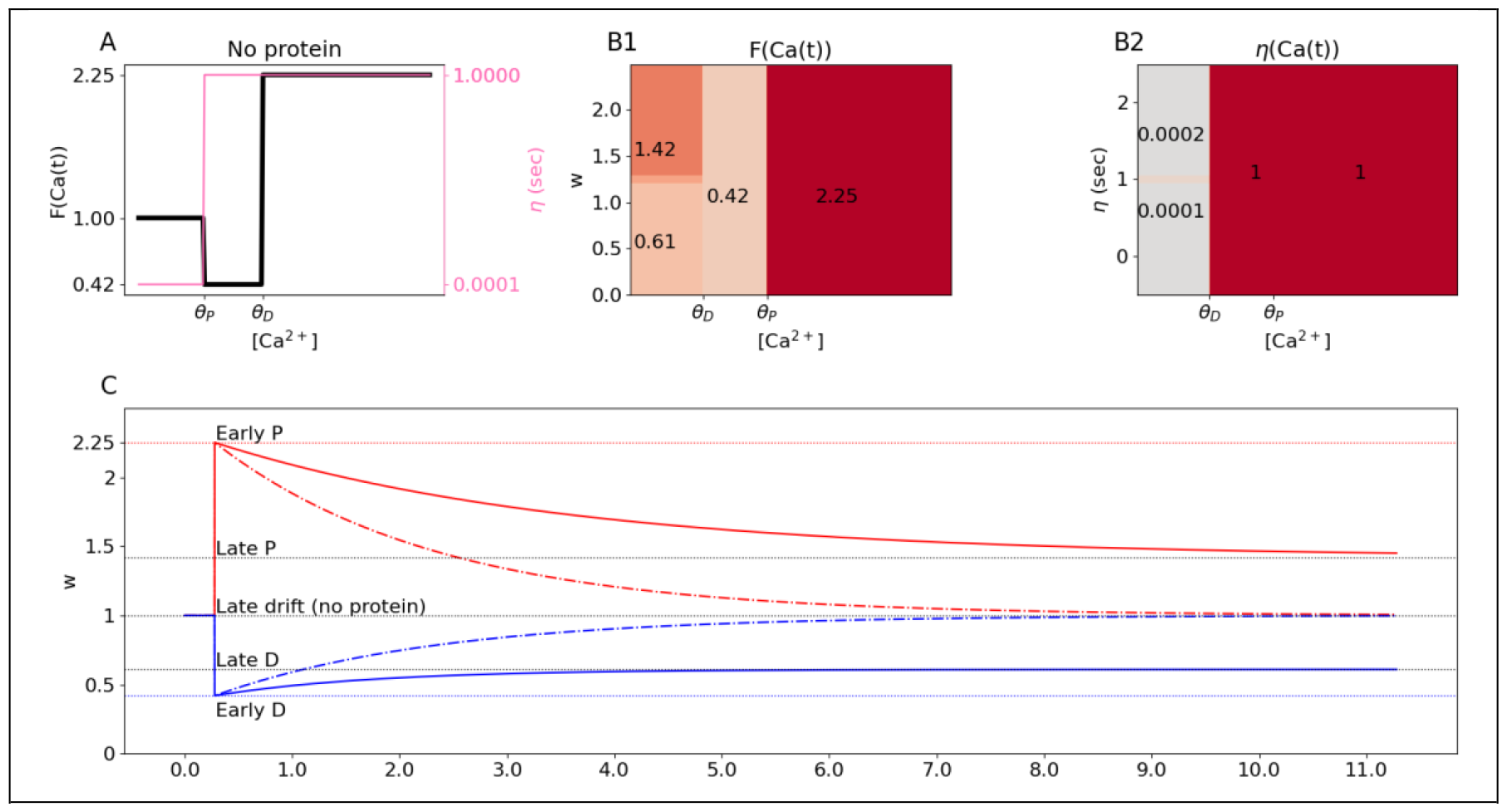
Protein-Dependent Late Phase Plasticity. **(A)** In the absence of protein synthesis, fixed points and learning rates can be described by one-dimensional functions of the [*Ca*^2+^], as in the one-dimensional FPLR rule (Fig. 1F-G). Here we use biologically realistic values for the late phase of plasticity (described in **Methods**). **(B)** If there are proteins to stabilize the new synaptic weights, fixed points (B1) and learning rates (B2) can be described as two-dimensional functions of [*Ca*^2+^] and the current weight *w*, as in the two-dimensional FPLR rule (Fig.4D). **(C)** Weights over time using the biologically realistic parameters from A and B in the presence (solid line) and absence (dotted line) for potentiative (red) and depressive (blue) levels of [*Ca*^2+^].

Where τ is the overall time constant of synaptic change and *γ*_*P*_ and *γ*_*D*_ are parameters that denote the rate of potentiation and depression, respectively.

The first term in Equation 3.1 is always active and expresses the calcium-independent dynamics of the slow drift to the UP or DOWN state given the current value of *ρ*. The second term in this equation is only active while the [Ca^2+^] is above the potentiation threshold and describes the asymptotic potentiative dynamics, while the final term is active in both the depressive and potentiative regions of [Ca^2+^] and describes the asymptotic depressive dynamics.

Eq. 3.1 can also be expressed as a step function (ignoring the noise term):

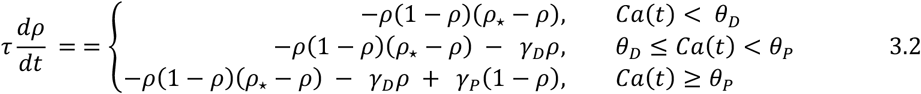

Note that in the depressive region of [Ca^2+^] (second line), the calcium-independent term (first term) continues to affect the synaptic efficacy based on the current value of *ρ*. In the potentiating region of [Ca^2+^] (third line), both the calcium-independent term and the depressive term are active in addition to the potentiating term.

### Simplified Graupner and Brunel Model

While the GB rule can describe a variety of experimental results, its dynamics can be complicated by the fact that multiple processes are active simultaneously – that is, the slow calcium-independent drift of the first term is always active, and the depressive process is always active when the potentiation process is active. From a modeling standpoint, this aspect of the GB rule may not be desirable, as the dynamics in each region of [Ca^2+^] do not exhibit simple asymptotic behavior, the fixed point for potentiation is not trivial to specify (note that *ρ* = 1 is not a fixed point of *ρ* if *Ca*(*t*) > *θ*_*P*_, see Fig. 3C2), and specifying *γ*_*P*_ is insufficient to know the actual rate of potentiation because the depressive term *γ*_*D*_ also affects the potentiation rate.

From a biological standpoint, it is also questionable whether depressive and potentiating processes are active simultaneously. While some studies have shown that depressive and potentiative mechanisms are operative at the same time and compete with each other (Burrell & Li, 2008; O’Connor et al., 2005b), another study (Cho et al., 2001) argues that once the [Ca^2+^] reaches the potentiation threshold, the depressive mechanisms are turned off. The slow bistable drift mechanisms in the first term of Eq 3.2 may also not be perpetually active; the long-term stabilization mechanisms required for late LTP/LTD (L-LTP and L-LTD) have been shown to be protein synthesis dependent, and this protein synthesis may only occur after the induction of early-LTP/LTD (Barco et al., 2008; Frey & Morris, 1997; Redondo et al., 2010).

We therefore propose a simplified version of the Graupner-Brunel rule which only has a single term active for any given concentration of calcium and synaptic efficacy value, resulting in a rule which achieves a qualitatively similar result but with more straightforward dynamics:

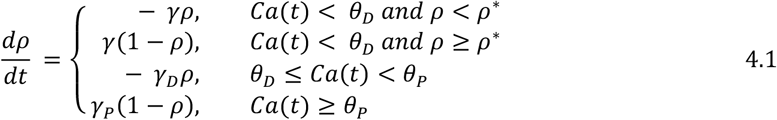

In this rule, in the pre-depressive region of [*Ca*^2+^], *ρ* drifts toward 0 at a rate of *γ* if *ρ* is below or equal to *ρ*^∗^(first line, note that there is now no unstable fixed point at *ρ*^∗^), or asymptotically toward 1 at a rate of *γ* if *ρ* is above *ρ*^∗^ (second line).In the depressive region of calcium (third line), *ρ* trends asymptotically toward the fixed point of 0 at a rate of *γ*_*D*_, and in the potentiative region of calcium (fourth line), *ρ* trends asymptotically toward the fixed point of 1 at a rate of *γ*_*P*_.

Unlike the original rule, the UP and DOWN states in the simplified rule occur precisely at *ρ* = 1 and *ρ* = 0 in all calcium regimes. Moreover, instead of a hyperbolic calcium-independent term, we use two linear terms to create a simple asymptotic increase or decrease toward the fixed points instead of a sigmoidal trend (Fig.3 - orange lines).

Additionally, the rate of change within each region of the [*Ca*^2+^] is specified by its own learning rate (*γ, γ*_*D*_, *γ*_*P*_). (We note that if one wishes to simulate the dynamics of simultaneously active depressive and potentiative mechanisms, this is still possible within our framework by designing a soft-threshold learning rate that varies gradually with the [*Ca*^2+^] to mimic the observed dynamics of the weight changes, as in Figure 2C. This illustrates the difference between the phenomenological and the mechanistic perspective; the phenomenological approach is agnostic as to whether depressive and potentiative mechanisms are competing, all that matters is the observed endpoint and rate of the resultant plastic changes for a given [*Ca*^2+^].)

Astute readers may notice that the simplified GB rule is similar in structure to the modified SBC rule. In fact, the second two lines of Eq 4.1 are identical to the modified SBC rule with fixed points {*F*(*θ*_*D*_ ≤ *Ca*(*t*) ≤ *θ*_*P*_) = 0, *F*(*Ca*(*t*) > *θ*_*P*_) = 1)} and learning rates {*η*(*θ*_*D*_ ≤ *Ca*(*t*) ≤ *θ*_*P*_) = *γ*_*D*_, *η*(*Ca*(*t*) > *θ*_*P*_) = *γ*_*P*_)}. The first two lines of Eq. 4.1, however, add a new feature, namely the dependence of the fixed points on the current weight *ρ*, in addition to the [Ca^2+^] (Fig. 4B).

### FPLR version of the GB rule

It is possible to generalize the simplified GB rule into a fully generic two-dimensional FPLR plasticity rule that specifies the fixed points and learning rates as a function of both the synaptic [Ca^2+^] and the current weight. Similar to the SBC rule, we have:

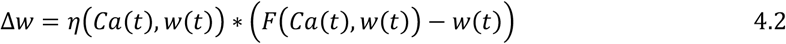

(We use *w* instead of *ρ* to specify the weight for consistency with the one-dimensional rule). Here, both the learning rates *η* and the fixed points *F* are two-dimensional step functions of both the [Ca^2+^] and the current weight *w*, as opposed to a one-dimensional step function of only the [Ca^2+^] in the SBC rule.

In addition to the bi-stable drift in the pre-depressive region of [Ca^2+^], Eq. 4.2 allows us to specify arbitrary numbers of weight-dependent fixed points in each region of [Ca^2+^].

However, when specifying fixed points of the weights as a function of the present weights, care must be taken to avoid overlapping basins of attraction. For example, if, in some region of [Ca^2+^], the fixed point for a synapse with a weight of *w* = 0.8 is *w* = 1, the fixed point for weight *w* = 0.9 must also be *w* = 1, because *w* = 0.8 must pass *w* = 0.9 on its way to *w* = 1.

One way to enforce this constraint is to specify *N* fixed points and *N* + 1 boundaries of the basins of attraction within each region of [Ca^2+^] such that the fixed points are always inside the closest basin boundaries on either side (Fig. 4C). For example, we can consider a rule that incorporates a tri-stable pre-depressive drift, where strong synapses drift to an UP state of *w* = 0.9, weak synapses drift to a DOWN state of *w* = 0.2, and synapses which aren’t particularly strong or weak drift toward a MIDDLE state of *w* = 0.5 (instead of an unstable fixed point of *w* = 0.5). (We intentionally choose fixed points in the pre-depressive region of [Ca^2+^] that are different from those in the potentiative (*w* = 1) and depressive (*w* = 0) regions of [Ca^2+^] to reflect experimental results that synaptic weights during the early phase of LTP and LTD can overshoot/undershoot the eventual weights to which they are stabilized (Manahan-Vaughan et al., 2000; Redondo et al., 2010).) We can specify the fixed points for the pre-depressive calcium region *F*(*Ca*(*t*) < *θ*_*D*_, *w*(*t*)) by choosing the boundaries of the basins of attraction as [0, 0.3, 0.7, ∞] and the fixed points as [0.2, 0.5, 0.9].

This translates into

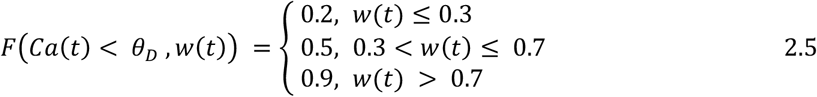

(Fig. 4C, 4D1-D4).

We can illustrate the dynamics of the tri-stable FPLR rule by applying our canonical protocol of a step of potentiative or depressive calcium for either a short or long duration. A synapse exposed to the short-duration stimulus will briefly potentiate or depress, but insufficiently to escape the MIDDLE fixed point’s basin of attraction, so it drifts back to the MIDDLE position. A synapse exposed long-duration stimulus, however, will escape the MIDDLE fixed point’s basin of attraction and thus drift to either the UP or DOWN fixed point after the calcium step is turned off (Figure 4D5).

### Incorporating protein-dependent late LTP/LTD

One of the major differences between the original SBC rule and the GB rule pertains to what happens in the “late phase” of LTP/LTD, hours after an LTP/LTD stimulation protocol is completed and [Ca^2+^] levels have returned to baseline. In the SBC rule the weight eventually drifts to 0 (or some other baseline state in the modified SBC rule), while in the GB rule weights are slowly stabilized to one of two stable points, an UP state or a DOWN state, depending on the synaptic strength at the completion of the plasticity protocol.

Biologically, the SBC rule is more representative of what has been observed in the absence of protein synthesis, and GB rule can be seen to correspond to the situations where proteins are synthesized to stabilize synaptic strengths for a longer period. In hippocampal cells, late-phase stabilization of potentiated and depressed synaptic states requires synthesis of proteins, without which synapses eventually drift back to their original strengths (Frey & Morris, 1997; Kauderer & Kandel, 2000; Redondo et al., 2010; Sajikumar et al., 2005). A similar phenomenon has been observed for LTD in Purkinje cells (Linden, 1996). This dependence on protein synthesis can be incorporated into the generic GB model by adding another dimension to the step functions for the weights and fixed points.

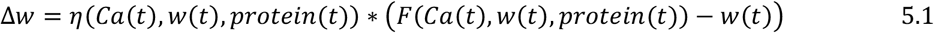

For illustration, if we make a simplifying assumption that there is a single stabilizing protein which can be either present (1) or absent (0), we might have the following rule:

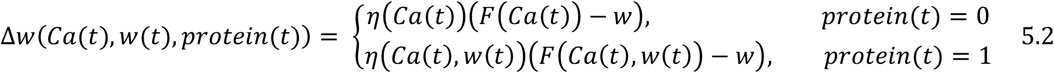

In other words, if there are no proteins to stabilize the new synaptic weights after a plasticity-inducing calcium stimulus, then synaptic weights simply drift to baseline, and the plasticity dynamics in each region of [*Ca*^2+^], including the pre-depressive region, can thus be described using a single weight-independent fixed point and learning rate in each region of [*Ca*^2+^], as in the generic SBC rule. However, if there are proteins to stabilize the new synaptic weights after a plasticity-inducing calcium stimulus, then fixed points and learning rates in the pre-depressive region of [*Ca*^2+^] are weight-dependent and the plasticity dynamics must be described with two-dimensional step functions, as in the generic GB rule. We illustrate these protein-dependent late phase dynamics using biologically realistic fixed points and time constants in Figure 5 (see **Methods**). Because the late phase dynamics operate on the order of hours and the early phase dynamics operate on the order of milliseconds, we can assume for simplicity that the early phase stimulation causes an instantaneous jump to the depressive or potentiative fixed point, i.e. we can set *η* = 1 *second* for the potentiative and depressive regions, while the late phase drift or stabilization occurs 4 orders of magnitude more slowly.

### Modeling frequency and spike timing dependent plasticity

Although the FPLR framework makes the dynamics of calcium-based plasticity more straightforward, it is still quite similar to the original SBC and GB rules, and one would not expect to observe substantial discrepancies in the implementation of plasticity inducing protocols. Nevertheless, for the sake of completeness, we reproduce two classic protocols: frequency-dependent plasticity and spike timing dependent plasticity (STDP). Most of the literature about these protocols pertains to the early phase of plasticity, and we therefore use the version of the one-dimensional FPLR rule where the pre-depressive drift is turned off by setting *η*(*Ca* < *θ*_*D*_) = 0, as in Fig. 1I.

To model the calcium itself, we use the simplified formalism (Graupner & Brunel, 2012) where calcium concentration at spine *k* is modeled as a value which jumps by *C*_*pre*_ whenever there is a presynaptic spike at spine *k*, jumps by *C*_*post*_ whenever there is a postsynaptic spike, and decays toward 0 at the rate of τ_*Ca*_ (Graupner & Brunel, 2012). Formally, this can be expressed by the equation:

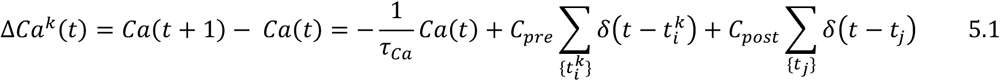

Where 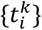 are the times of the presynaptic input spikes at spine *k* and {*t*_*j*_} are the times of the post synaptic spikes.

We first demonstrate how calcium-based plasticity results in synaptic changes that depend on the frequency of the presynaptic input. Experimentally, low-frequency stimulation (LFS) tends to produce depression, while high-frequency stimulation (HFS) tends to produce potentiation (O’Connor et al., 2005b). Intuitively, if *C*_*pre*_ is below *θ*_*D*_, a single presynaptic input spike will not produce depression, but if several input spikes occur such that each spike rides on the tail of the previous spike, the [*Ca*^2+^] can build up such that it rises above *θ*_*D*_, causing depression, and if the spikes occur with sufficiently high frequency, the [*Ca*^2+^] can rise above *θ*_*P*_, resulting in potentiation (Figure 6A).

**Figure 6:**
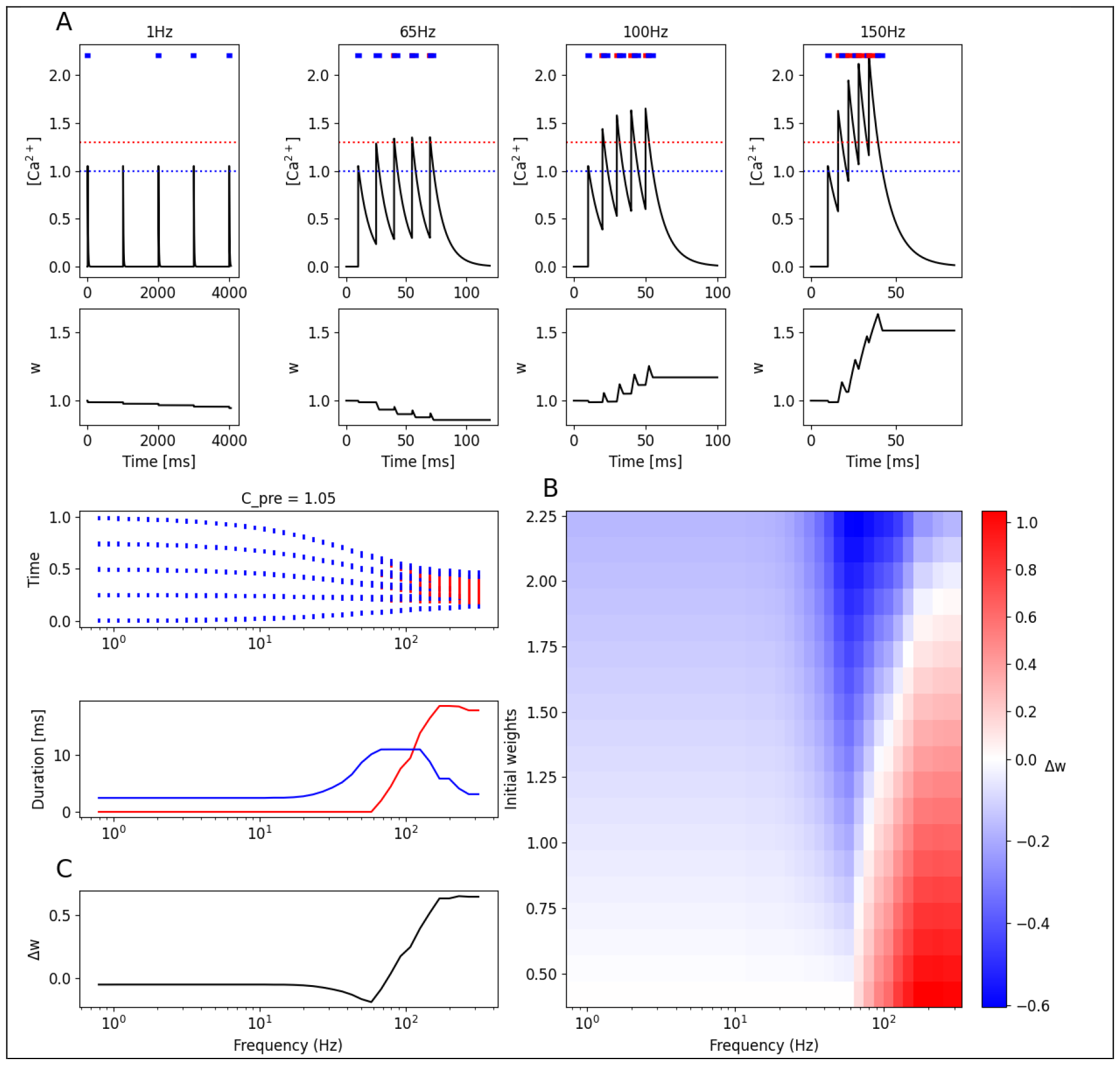
Frequency-Dependent Plasticity. **(A)** Calcium (top) and weight changes (bottom) produced by a train of 5 presynaptic inputs at different frequencies. Dashed blue and red lines indicate *θ*_*D*_ and *θ*_*P*_, respectively. Blue and red stripes at the top of the plots form a bar code indicating when times at which the calcium is in the depressive (blue) or potentiative regions (red). **(B)** (Top) Bar codes (As in A1) oriented vertically as a function of input frequency (ISI). Time axis is normalized to beginning and end of each simulation. (Middle) Total duration in depressive (blue) or potentiative (red) regions of [*Ca*^2+^] as a function of input frequency. (Bottom) Total change in weight (final-initial) for each input frequency. **(C)** Heatmap of plastic changes as a function of both initial weight and input frequency. Note that lower weights are easier to potentiate, higher weights are easier to depress.

In order to determine the final synaptic weight after a plasticity protocol, it is not sufficient to simply compare the duration above *θ*_*D*_ and *θ*_*P*_ due to the difference in rates of the depressive and potentiative processes as well as the asymptotic dynamics of the plasticity, which make it easier to depress large weights and potentiate small weights. It can therefore be helpful to create a “bar code” for different stimulation protocols by indicating the times at which the [*Ca*^2+^] was above the *θ*_*D*_ and *θ*_*P*_, which can help give a more intuitive feel for which protocols will result in synaptic potentiation or depression (Figure 6A-B).

Because the synaptic changes in the FPLR framework occur in an asymptotic manner, it is easier to potentiate weak synapses and easier to depress strong synapses. This means that the result of a frequency-dependent plasticity protocol will depend on the initial synaptic weight. If a synapse starts out closer to the depressive fixed point, it is easier to potentiate, while if it starts out closer to the potentiative fixed point, it is easier to depress. In fact, synapses which start out with large weights may even be depressed by a high-frequency protocol (Figure 6B).

The reason for high-frequency protocols depressing strong synapses in this model is subtle: whenever the [*Ca*^2+^] rises past *θ*_*P*_, it will inevitably spend some time in the depressive region when the stimulation ends and the calcium decays to the baseline. Thus, for synaptic potentiation to be maintained, the magnitude of potentiation must be sufficiently large to not be completely erased by the subsequent depression. However, if the synapse starts out with a weight near the potentiation fixed point, virtually no potentiation can occur, so the depression during the decay phase of the calcium will be the only effect observed.

This “what goes up must come down” effect is an inevitable quirk of calcium threshold models which incorporate decaying calcium signals, and while this quirk can sometimes be helpful in explaining some experimental results, it creates complications for reproducing other experimental results, and it is also a bit counterintuitive. There is some experimental evidence that potentiation protocols will “lock in” potentiation to prevent subsequent depression (O’Connor et al., 2005b), which may help to alleviate this “what goes up must come down” problem. Our simulations here, however, do not include a lock-in feature, so potentiation protocols will always include a period of depression once the stimulation concludes and the [*Ca*^2+^] decays back to baseline. It is also possible to avoid this issue if the rate of depression is substantially slower than the rate of potentiation and the decay of the calcium is sufficiently fast such that the amount of depression that occurs while the calcium is decaying back to baseline after a potentiation protocol is negligible.

### Spike Timing-Dependent Plasticity (STDP)

We can use our intuition from the above section about frequency-dependent plasticity to build an FPLR-based simulation of spike-timing dependent plasticity (STDP). We wish to replicate the “classic” STDP curve, where presynaptic input before postsynaptic stimulation causes potentiation at the activated presynaptic synapse, whereas postsynaptic spiking before presynaptic input causes causes depression, and both effects decrease with increased time intervals (Bi & Poo, 1998).

To reproduce this result, we simulated the calcium generated by single presynaptic spike (which creates a [*Ca*^2+^] pulse of height *C*_*pre*_) at 100 ms of a 200 ms simulation. We then generate a postsynaptic spike (which creates a [*Ca*^2+^] pulse of height *C*_*post*_) either before or after the presynaptic spike at varying timing intervals. By appropriately setting the values for *C*_*pre*_ and *C*_*post*_, it is possible to replicate the classic STDP curve (different parameter values can result in different STDP curves, see (Graupner & Brunel, 2012)). As with the frequency-dependent protocol above, he magnitude of synaptic weight change in the STDP protocol will depend on the initial synaptic weight; large weights are faster to depress and small weights are faster to potentiate.

### Modeling Behavioral Time Scale Plasticity

Recent experimental findings in the hippocampus have revealed a novel form of plasticity, known as behavioral time scale plasticity (BTSP) (Bittner et al., 2015, 2017; Milstein et al., 2021). A mouse running on a treadmill can spontaneously form hippocampal place fields when the soma in injected with a strong current, inducing a plateau potential.

After a single induction, this plateau potential results in the neuron exhibiting a place field selective to the mouse’s location few seconds before or after the time of the plateau potential. Moreover, this place field can be modified; if a second plateau is induced while the mouse is at a different location near the first place field, the place field will shift to the new location, thus “overwriting” the first place field. However, if the second location is sufficiently far away from the neuron’s first place field, the neuron forms an additional place field at the second location without “overwriting” the first place field (Milstein et al., 2021).

Although BTSP has been previously modeled using an “eligibility trace” approach (Cone & Shouval, 2021; Gerstner et al., 2018; Milstein et al., 2021), we argue that the FPLR framework for calcium-based plasticity is sufficient to account for many of the results from the BTSP experiments. (See **Discussion** for further comments on the biological plausibility of each modeling approach.) The basic intuition for how calcium control can result in BTSP is similar to Hebbian plasticity. Neither the presynaptic input nor the plateau potential themselves bring enough calcium into a spine for a long enough time to induce substantial changes to the synaptic weight. When the [*Ca*^2+^] traces from both the presynaptic sensory input and plateau potential coincide (or the [*Ca*^2+^] released from internal stores in the endoplasmic reticulum induced by these events, see **Discussion**), the combined [*Ca*^2+^] from both sources rises above the potentiation threshold for around a second, inducing potentiation at the synapses that were active at the time of plateau induction. Because the potentiated sensory inputs all correspond to the mouse’s location at the time of plateau induction, these potentiated synapses comprise a place field at that location.

The reason why a second induction near to the location of the first induction will overwrite the first place field is because the partially decayed calcium trace from the plateau combines with the calcium from the presynaptic input corresponding to the first place field to surpass the depression threshold but not the potentiation threshold, depressing the synapses from the first place field. (This is if the second place field precedes the first place field on the track. If the first place field precedes the second place field on the track, the depression results from the decaying local calcium signal from the presynaptic input from the first place field combining with the [*Ca*^2+^] from the plateau potential.) If the two locations are far enough apart, however, the calcium from the plateau potential will have decayed sufficiently such that it will not surpass the depression threshold for a substantial amount of time when added to the local calcium from the presynaptic input (or vice versa). Finally, the reason why only synapses that were previously potentiated are depressed when “overwritten” in the BTSP protocol is because of the weight dependence of the FPLR rule. At baseline, all synapses that do not take part in a place field have weights near the depressive fixed point, so they cannot be further depressed.

To illustrate this, we simulated a mouse running at constant velocity on a circular track by sequentially presenting track locations as inputs to a leaky integrator model neuron with spatially-tuned presynaptic inputs. Calcium could enter a synapse as a consequence of its local input or due to the supervising signal, which induces a plateau potential in the neuron and globally broadcasts calcium to all synapses. The synaptic calcium decayed with a time constant of ∼2 seconds (Fig. 8, see **Discussion** regarding the question of biologically plausible calcium decay time constants and potential implementation involving internal calcium stores). The synaptic weights of the model neuron implemented the FPLR calcium-based plasticity rule described in Equation 2.1 with the drift rate set to 0 as in the previous simulations.

**Figure 7:**
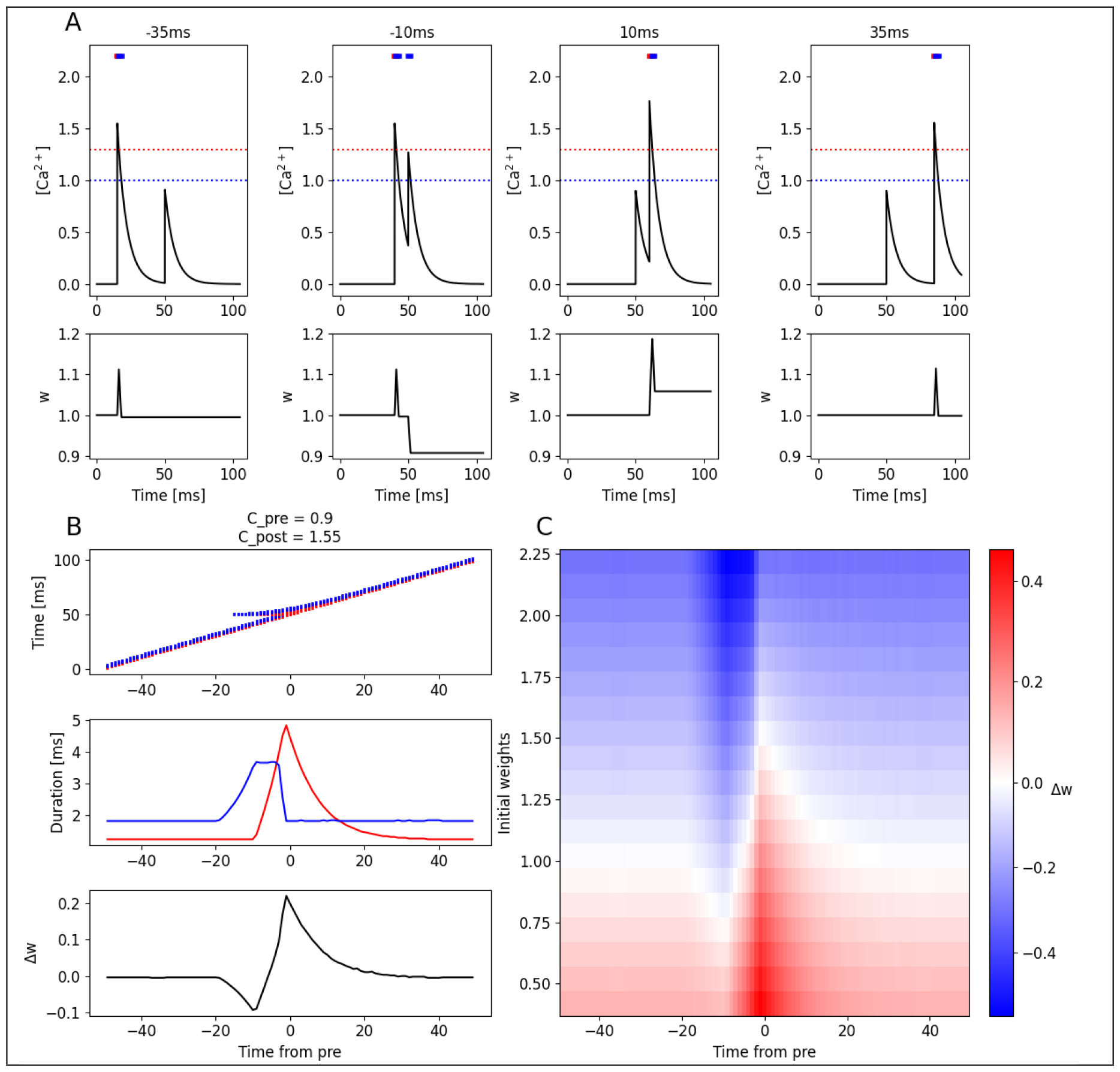
Spike Timing Dependent Plasticity (STDP) **(A)** Calcium (Top) and weight changes (bottom) produced by a single presynaptic and postsynaptic stimulation at different time intervals. Negative time interval indicates post-before-pre, positive time interval indicates pre-before-post. **(B)** Bar codes (Top), durations (Middle), and weight changes (Bottom) as a function of post-pre interval. **(C)** STDP heatmap as a function of post-pre interval and initial weights.

**Figure 8:**
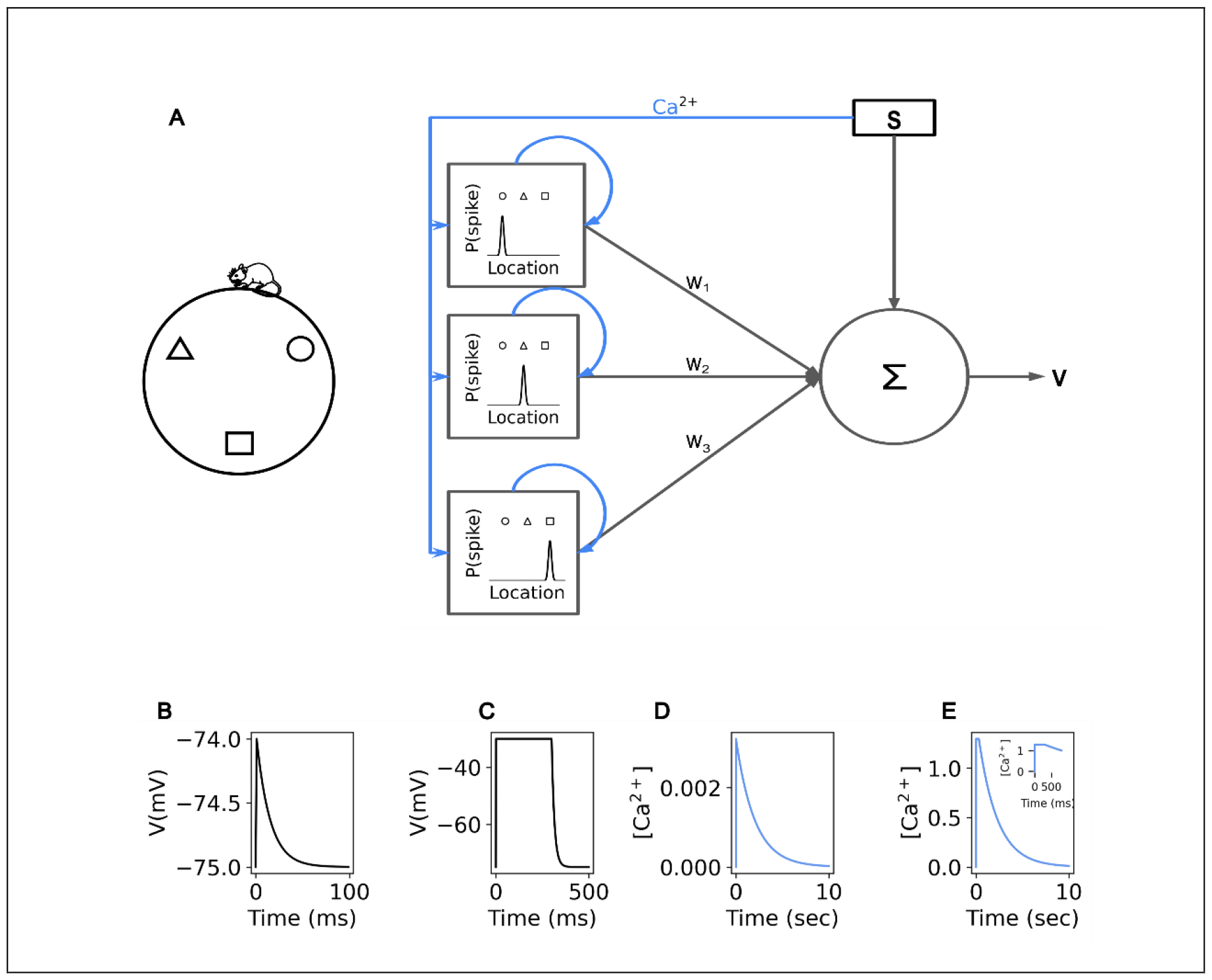
Simulation and Neuron Model for Behavioral Time Scale Plasticity. **(A)** (Left) Schematic of a mouse running along a circular track. Shapes indicate position along the track. (Right) A leaky integrator neuron sums weighted current input (black arrows) from each of its spatially-tuned synapses (squares). Spatial input also induces calcium (blue arrows) locally at each synapse. A supervising signal (S) can induce a plateau potential in the postsynaptic neuron as well as globally broadcast calcium to all synapses simultaneously. **(B)** Voltage response of the neuron to a single presynaptic spike. **(C)** Voltage response of the neuron to a supervisor-induced plateau potential. **(D)** Calcium response of a synapse to a single presynaptic spike at that synapse. **(E)** Calcium response at one synapse to a supervisor-induced plateau potential. Inset: zoom-in on the first 800 milliseconds to show calcium plateau.

The track was presented seven times to the neuron, simulating seven laps, with each lap taking ten seconds to run. At every location on the track, presynaptic inputs whose receptive fields overlapped with that location would contribute current to the neuron as well as induce a calcium signal with height *C*_*pre*_ at the associated “postsynaptic spine”. The presynaptic calcium signal decayed exponentially at a rate of τ_*pre*_. On the first lap, no induction was performed, establishing a baseline of activity in the absence of a place field.

During the second lap, a plateau potential was induced at 3.5 seconds into the lap depolarizing the neuron’s voltage and inducing a step of calcium for 300 milliseconds with height *C*_*plateau*_ at all synapses. After the plateau induction, the calcium from the plateau decayed exponentially at a rate of τ_*post*_ . During the third lap, a voltage ramp was observed from ∼ 1.5-5 seconds into the lap indicating that the plateau induction from the previous lap had produced a place field. In the fourth lap, a plateau potential was induced at 2 seconds into the lap. During the fifth lap, a voltage ramp was observed from ∼ 0-3.5 seconds into the lap indicating that the preciously induced place field had been “overwritten” by the plateau induction in the fourth lap. In the sixth lap, a plateau potential was induced at 7.5 seconds into the lap. During the seventh lap, two voltage ramps were observed, one ramp from ∼ 0-3.5 seconds and the other from ∼ 5.5-9 seconds into the lap indicating that the plateau from the sixth lap induced a new place field but the place field observed at the fifth lap was not erased (Fig. 9).

**Figure 9:**
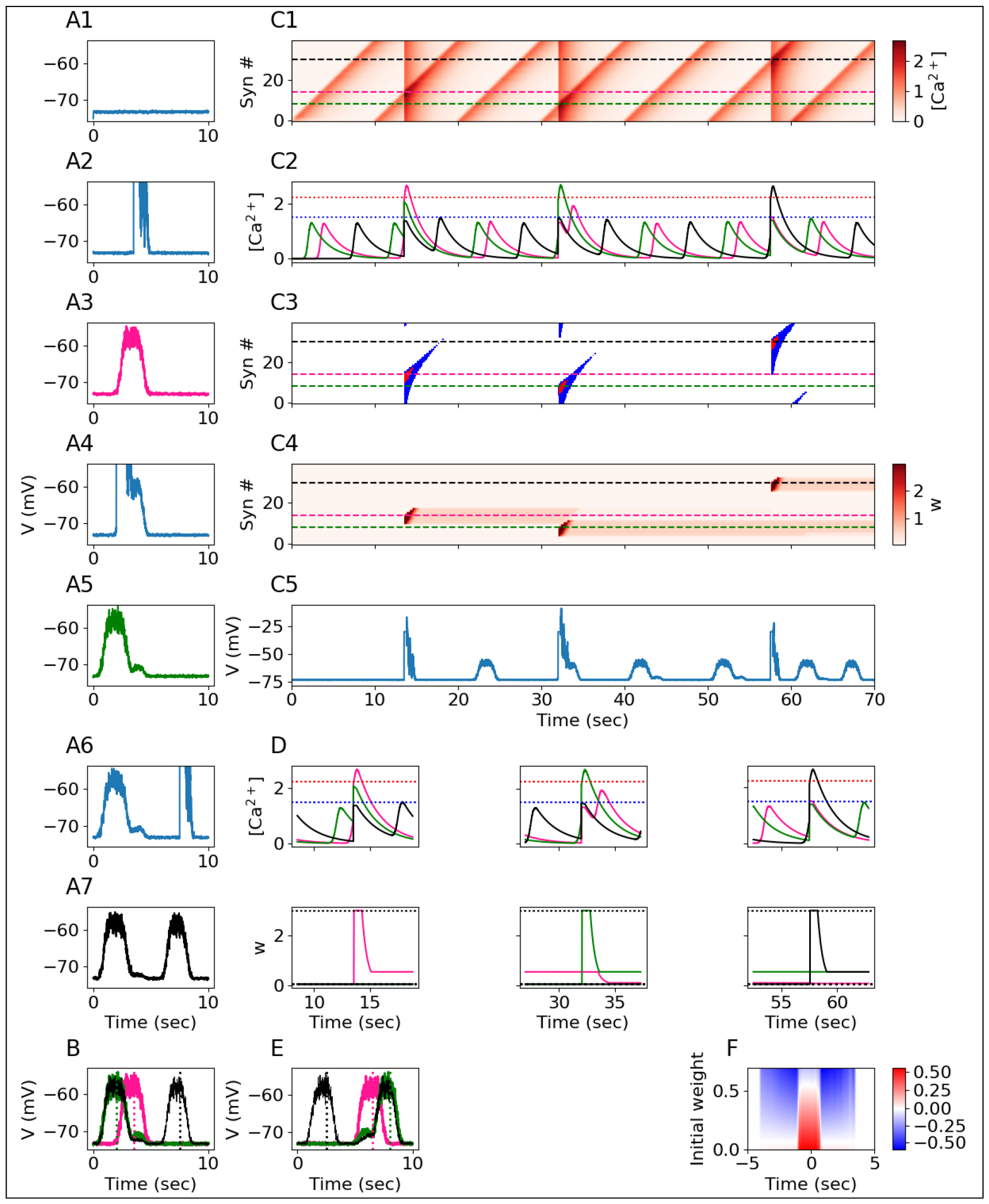
Behavioral Time Scale Plasticity. **(A1-A7)** Voltage traces for 7 laps, each lap lasts for 10 seconds. A plateau potential is induced in the 2nd, 4th, and 6th laps. Green, pink, and black traces indicate place fields observed on the 3rd, 5th, and 7th laps, respectively. **(B)** Overlay of voltage traces for place fields from the 3^rd^ (pink), 5^th^ (green), and 7^th^ (black) laps. Vertical dotted lines indicate plateau induction location during the preceding lap. **(C1)** Total [*Ca*^2+^] per synapse (rows, 40 synapses) over the course of all 7 laps. Dashed lines indicate three synapses whose receptive fields are centered at the location of the first (pink, S3.5, see text) second (green, S2.0) or third (black, S7.5) plateau induction. **(C2)** [*Ca*^2+^] traces for the three synapses indicated by the horizontal lines in (C1). Blue and red dotted lines indicate *θ*_*D*_ and *θ*_*P*_. **(C3)** Plasticity bar codes for each synapse (rows) over all laps. **(C4)** Weights over time for each synapse over all laps. **(C5)** Voltage over time, as in A, for all 7 laps. **(D)** Zoom-in on the calcium traces (top) and weights (bottom) from the three synapses shown in B2 at the time of each plateau induction. In the second lap (left), the pink synapse’s [*Ca*^2+^] rises beyond *θ*_*P*_, inducing potentiation toward the maximum strength (dotted horizontal line at top), and the green synapse’s [*Ca*^2+^] rises above *θ*_*D*_, although the green synapse is already at the minimum strength (dotted horizontal line on bottom) so it can’t depress any further. In the fourth lap (middle), the green synapse’s [*Ca*^2+^] rises beyond *θ*_*P*_, inducing potentiation, and the pink and black synapses’ [*Ca*^2+^] rises above *θ*_*D*_, but only the pink synapse can be depressed because it was previously potentiated. In the sixth lap (right), the black synapse’s [*Ca*^2+^] rises beyond *θ*_*P*_, inducing potentiation. **(E)** Result of a “mirror image” experiment, where the locations of plateau inductions are reversed (see supplementary figure S1). Green, pink, and black traces show laps 3,5, and 7; dotted vertical lines indicate induction locations from laps, 2,4, and 6, as in (B). **(F)** Change in weights as a function of initial weight and receptive field distance from plateau onset.

To show how the FPLR calcium rule produced these results we visualized the [*Ca*^2+^], barcodes, and weights of each of the synapses over the entire course of the experiment (Fig. 9C) focusing on the three synapses whose place fields were centered around the track location of the plateau inductions of laps 2, 4 and 6 (Fig. 9B2, 8C). We call these synapses S3.5 (pink trace), S2.0 (green trace) and S7.5 (black trace), respectively, according to the track location (in units of seconds from beginning of track) at the center of their receptive fields. At the time of the first plateau induction, the [*Ca*^2+^] for S3.5 surpasses the potentiation threshold for ∼ 1 second, potentiating it (and synapses with nearby receptive fields). Because of the “what goes up must come down” effect, S3.5 depresses substantially after it is potentiated due to the [*Ca*^2+^] spending some time in the depressive region while it decays.

However, S3.5 still remains somewhat potentiated. The [*Ca*^2+^] for S2.0 also enters the depressive region, however because all the synapses were initialized at the depressive fixed point, S2.0 cannot be further depressed. This illustrates how the weight dependence of the FPLR rule has important functional consequences for place field formation. Without the weight dependence, there would always be a depressed region around the potentiated location, resulting in a Mexican hat-shaped place fields. The weight dependence ensures that only previously-potentiated synapses are ever depressed.

When the second plateau potential is induced in the fourth lap, the [*Ca*^2+^] for S2.0 surpasses the potentiation threshold for ∼ 1 second, potentiating it (with some subsequent depression) as before. The [*Ca*^2+^] of S3.5 enters the depressive region for ∼ 1 second due to the combination of the local presynaptic [*Ca*^2+^] and the partial [*Ca*^2+^] from the decayed plateau potential. This results in the depression of S3.5 back to the depressive fixed point was previously. The weight S7.5 does not change, as before.

Finally, at the time of the third plateau induction, the [*Ca*^2+^] at S7.5 surpasses the potentiation threshold for ∼ 1 second, potentiating it. Because the plateau induction at 7.5 seconds into the track was sufficiently temporally distanced from the activation of the receptive fields of S2.0 and S3.5, the decayed local and supervisory calcium signals did not overlap to surpass the depression threshold for a significant amount of time, resulting in only a negligible magnitude of depression for S2.0 (Fig. 9C-D).

To ensure that all the above results hold when the locations of the inductions are reversed, we performed a “mirror image” experiment, where plateau inductions at laps 2, 4. and 6 occurred at 6.5 seconds, 8 seconds, and 2.5 seconds from the beginning of the track, respectively. The mirror image experiment yielded similar results (Fig. 9E, Fig S1).

To fully characterize the expected weight change induced by a plateau potential for synapses with different initial weights and receptive fields, we simulated a mouse running a single lap with a plateau potential induced in the middle of the lap (at 5 seconds). Using the [*Ca*^2+^] at each synapse we calculated the magnitude and direction of plasticity at each synapse for a range of initial weights. Synapses with receptive fields selective to locations near the center of the track potentiate if they started out with low weights but depress if initialized with large weights (due to the fact that synapses with large weights can’t potentiate much, but do depress when the calcium signal decays because of the “what goes up must come down” effect). Synapses with receptive fields far from the center of the track will depress slightly if they are initialized with large weights but will not appreciably change if initialized with low weights (Fig. 9E). This profile is qualitatively consistent with the experimental results from (Milstein et al., 2021).

## Discussion

In this work, we have developed a straightforward mathematical framework, the FPLR rule, to describe calcium-dependent long-term plasticity dynamics. The FPLR framework, based on the rules of (Graupner & Brunel, 2012; Shouval et al., 2002), enable modelers to describe plasticity dynamics in each region of [*Ca*^2+^] by specifying fixed points and learning rates as one-dimensional step functions of the [*Ca*^2+^] or as two-dimensional functions of both the [*Ca*^2+^] and the current value of the weight. This makes it simple to model novel experimental results such as the examples of Purkinje neurons and additional no-plasticity zones that we showed above. Additionally, we have shown the one-dimensional and two dimensional can be integrated into a single framework wherein SBC-like decay dynamics occur in the late phase of plasticity in the absence of protein synthesis, whereas GB-like stabilization dynamics occur in the late phase when proteins are available to stabilize weight changes made in the early phase of plasticity. The FPLR framework, which allows for an arbitrary number of fixed points, also fits well with the theoretical and experimental literature suggesting that synaptic weights have on the order of 10 discrete states comprised of nanoclusters of AMPA receptors (Bartol et al., 2015; Liu et al., 2017).

To demonstrate the ability of the FPLR framework to reproduce classic plasticity protocols, we used the FPLR approach to implement frequency-dependent and spike timing dependent plasticity. We introduced the technique of plasticity “bar codes” as a simple way of keeping track of the times when the [*Ca*^2+^] was in the potentiative or depressive region. We showed that due to the saturating nature of plasticity, the outcome of frequency-dependent plasticity and STDP (indeed, any plasticity protocol) depends on the initial synaptic weights–it is easier to depress strong weights and potentiate weak weights. Finally, we showed that this weight-dependence of the FPLR framework enables us to explain a novel experimental result – behavioral time scale plasticity (BTSP).

### Alternative Calcium-Based Plasticity Models

We note that both the SBC and GB rules as well as the FPLR framework assume that synaptic weight changes depend on the magnitude of [*Ca*^2+^]. However, there are other theories as to how plasticity might depend on [*Ca*^2+^]. For example, changes in synaptic weights may depend both on the duration and magnitude of the calcium signal in a way that is not captured by the SBC and GB rules. Alternatively, the source of calcium – i.e. NMDA receptors, voltage-gated calcium channels or internal calcium stores – may affect long term plasticity due to different second messengers being localized to specific “nanodomains” near the different calcium sources (For a review of alternative models, see (Evans & Blackwell, 2015)).

More detailed models have also been developed that explicitly incorporate calcineurin and CamKII activity (Li et al., 2023; Rodrigues et al., 2021) as well as other presynaptic and postsynaptic mechanisms (Ebner et al., 2019). It is also possible to explicitly model protein dynamics for plasticity stabilization, for example via synaptic tag-and-capture model (Clopath et al., 2008). While more detailed plasticity rules may enable more specific experimental predictions, over 100 molecules have been implicated in long-term plasticity (Sanes & Lichtman, 1999), making it effectively impossible to have truly comprehensive mechanistic model. Our phenomenological framework can thus be useful for creating models that have relatively few parameters while still capturing essential aspects of calcium-based plasticity. Our fixed point–learning rate approach can also be extended to incorporate other molecular mechanisms, as we suggested with protein synthesis-dependent late-phase plasticity, although some molecular processes may not be well-described in this framework.

Additional model complexity may also be needed to model plasticity in physiological concentrations of extracellular calcium (Inglebert et al., 2020; Inglebert & Debanne, 2021).

### Modeling Calcium

Throughout this work, we have used simple exponentially decaying calcium stimuli to demonstrate the dynamics of our plasticity rules. When modeling neurons in a detailed fashion, it will usually be necessary to explicitly model the calcium influx from various sources, such as voltage-gated calcium channels and NMDA receptors, in order to explore the plastic consequences of presynaptic and postsynaptic activity. While detailed modeling of calcium dynamics is beyond the scope of this paper, work in this direction can be found in the original SBC and GB papers (Graupner & Brunel, 2012; Shouval et al., 2002) as well as elsewhere in the modeling literature (Chindemi et al., 2020).

### Modeling Behavioral Time Scale Plasticity

As we noted in **Results**, previous attempts to model BTSP suggested an “eligibility trace” approach, where separate presynaptic and postsynaptic signals must interact with each other to produce plasticity (Cone & Shouval, 2021; Gerstner et al., 2018; Milstein et al., 2021). However, given the extensive evidence for the calcium basis of plasticity in other contexts, and the fact that NMDA and VGCC channel blockers disrupt BTSP, (Bittner et al., 2017) we argue that it is worthwhile to give serious consideration to the possibility that calcium is the mechanism underlying BTSP.

One reason why eligibility traces were used in previous work (Cone & Shouval, 2021) is that classical calcium-based protocols like STDP occur at timescales that are faster than would be necessary to model BTSP. Indeed, in our own replications of STDP and frequency-dependent plasticity, we used time constants for the decay of calcium on the order of 10 milliseconds, while modeling BTSP required calcium decay time constants on the order of seconds. Calcium imaging data often shows calcium decay constants on the order of hundreds of milliseconds and that decay constants can vary widely due to various active and passive mechanisms (Majewska et al., 2000). However, time constants extracted from calcium imaging may not necessarily be reliable for modeling calcium-based plasticity, as the calcium indicator itself may interfere with the dynamics of the calcium. Indeed, more recent work has shown that calcium indicators with lower binding affinity show smaller decay constants, on the order of tens of milliseconds (Miyazaki & Ross, 2022).

We suggest that various molecular mechanisms inside the cell, such as the manipulation of calcium pumps, or intracellular calcium induced calcium release (CICR) from internal calcium stores (Rose & Konnerth, 2001), can titrate the speed at which calcium decays, enabling calcium-based plasticity to operate on a spectrum of timescales. Recent experimental and modeling studies have also shown the importance of intracellular calcium release for BTSP (Caya-Bissonnette et al., 2023; O’Hare et al., 2022). Although the free calcium observed via calcium imaging decays at fast time scales, presynaptic or postsynaptic events may trigger influx of calcium into intracellular calcium stores in the endoplasmic reticulum, which is later released. As such, the calcium traces in our model can represent a simplified description of the net [*Ca*^2+^] contained both in the cytosol and the intracellular calcium stores, the calcium from the latter being released to the cytosol when presynaptic stimulation or a supervising signal occurs. Such a process could be more precisely described by explicitly modeling CICR dynamics (Caya-Bissonnette et al., 2023). The dynamics of second messengers, such as CaMKII and phosphatases, may also underlie the longer timescales of BTSP (Jain et al., 2023; Li et al., 2023).

### Future Directions

The FPLR framework suggests a standardized experimental paradigm to characterize plasticity dynamics. Namely, [Ca^2+^] should be fixed at the potentiative or depressive value for a sufficient duration such that the synaptic weight no longer changes (or by performing an analogous LTP or LTD protocol such that weight saturation is observed). Then the synaptic strength should be observed over a period of hours or longer to observe the eventual late-phase drift fixed points. From these observations it is straightforward to characterize the FPLR fixed points (i.e. the weight saturation and drift values) as well as the learning rates by fitting a simple exponential to the experimentally observed dynamics.

Without performing these experiments, it is difficult to disambiguate the fixed points and the plasticity rates. A synapse which is potentiated via an HFS protocol can be potentiated to an even higher strength by performing another HFS protocol, thus merely looking at the peak EPSP value after a standard LTP/LTD protocol is insufficient to determine the potentiative or depressive fixed point (Enoki et al., 2009). Performing the necessary experiments to find fixed points and learning rates among different cell types, species, brain regions and even at different dendritic locations within the same neuron can lead to a deeper understanding of long-term plasticity.

In addition to enabling experimentalists to model their results, the FPLR framework is sufficiently simple and flexible that theoreticians can incorporate them into neuron models of varying levels of complexity, ranging from simple point neuron models to detailed biophysical models. This can open avenues toward understanding how calcium-based plasticity can lead to learning at the single neuron or even network-level resolution. The flexibility of our framework to specify arbitrary fixed points and plasticity rates can also enable the exploration of activity-dependent changes in plasticity rules, known as metaplasticity (Abraham, 2008).

## Methods

All figures were created in python using the Numpy (Harris et al., 2020) and Matplotlib (Hunter, 2007) packages.

### Plasticity parameter estimates

Everywhere in the paper except for Figure 5, learning rates and fixed points are meant to convey qualitative understanding of the dynamics of plasticity, not biologically realistic parameters.

For Figure 5, we used the following experimental results to approximate biologically realistic time constants: in rat hippocampus, for LTD (*in vivo*), in the early phase of depression, synapses depress to 42% of their initial strength after initializing a low-frequency stimulation (LFS) (900 pulses at 1 HZ) and either drift back to ∼92% of their baseline in the presence of anisomycin after 3-4.5 hours (which blocks protein synthesis) or stabilize at ∼61% of their baseline in the absence of anisomycin after 3-4.5 hours (Manahan-Vaughan et al., 2000). For LTP (*in vitro*), in the early phase of potentiation, synapses potentiate to ∼225% of their initial strength minutes after initializing a high-frequency stimulation protocol (100 pulses at 100 Hz, performed 3 times) high-frequency stimulation (HFS) and either drift back to baseline in the presence of anisomycin after 10 hours or stabilize at ∼142% of their baseline in the absence of anisomycin after 10 hours (Redondo et al., 2010).

To calculate the time constants, we assumed that the calcium signal from each pulse lasted ∼10 ms. As such, the calcium signal lasted ∼3 seconds and the LTD calcium signal lasted ∼9 seconds.

Because different time constants were found for the drift back to baseline for LTP and LTD in the presence of anisomycin, we used the mean of the learning rates that were calculated (0.0046) from both experiments, although in principle it is possible to use a 2-dimensional rule to differentiate the drifts down from the potentiated state or up from the depressed state even in the absence of protein synthesis.

The fixed points for the weights in Figure 5 were not precisely calibrated to experimental data due to substantially different baselines in peak EPSP measured in various plasticity experiments (Enoki et al., 2009; Manahan-Vaughan et al., 2000; Redondo et al., 2010). The values used for fixed points in Figure 5 can be thought of as “change relative to baseline” where value of 1 indicates no change. When modeling experimental data, depending on the context, “synaptic weights” may refer to AMPA conductance, number of AMPA receptors/nanoclusters, integral or peak EPSP or EPSC measured at the dendrite or the soma, spine head volume or area, number of docked vesicles, release probability, or other parameters; fixed point values can be expressed directly in units of the relevant parameters.

### Behavioral Time Scale Plasticity Simulation

To simulate behavioral time-scale plasticity (BTSP), we used a leaky integrator neuron where the voltage *V* is determined by the differential equation:

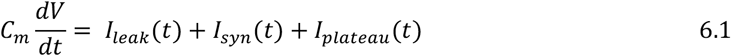

Where *C*_*m*_ = 1 nF, *I*_*leak*_ is the leak current, *I*_*syn*_ is the total contribution to the postsynaptic neuron from all presynaptic inputs, and *I*_*plateau*_ is the current contribution from a plateau potential induction. (As we are mainly interested in the subthreshold ramp activity rather than postsynaptic spikes, this model neglects postsynaptic spiking activity). The leak current is defined as:

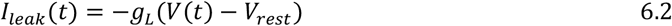

Where *g*_*L*_ is the leak conductance. *V*_*rest*_ is the resting potential of the membrane. It is helpful to define the voltage leak time constant 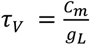. In the absence of input, the voltage thus decays according to the equation 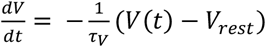

The presynaptic current is defined as:

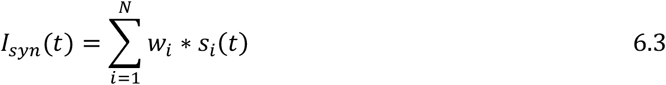

where *w*_*i*_ is the weight of synapse *i* and *s*_*i*_ (*t*) is a binary variable indicating whether synapse *i* produced a spike at time *t*. The probability of a presynaptic spike at synapse *i* was defined as:

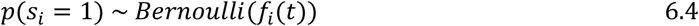

Where *f*_*i*_(*t*) is a bell-shaped receptive field centered around the neuron’s preferred track location *l*_*i*_. (We can define locations in terms of milliseconds of time from the start of the lap because we assume the mouse runs at constant velocity.) If the mouse’s location is given as *l*(*t*) = *t mod T*, where *T* is the track length in units of milliseconds of running time, we have:

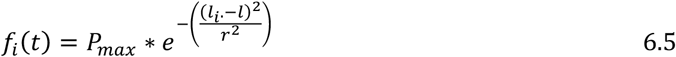

where *P*_*max*_ is the probability of the neuron firing a spike when the mouse is at the center of the receptive field of that neuron, and *r* determines receptive field width. The presynaptic receptive fields thus tile the track length with one receptive field center every 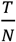 milliseconds of running distance.

To ensure that the plateau potential took the form of a rectangular voltage clamp step, we defined the plateau current as:

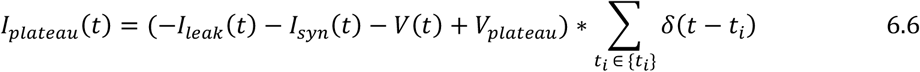

Where *V*_*plateau*_ is the target steady state voltage during the plateau induction and {*t*_*i*_} is the set of all times at which the plateau induction is active.

The [*Ca*^2+^] at each synapse *i, Ca*^*i*^, is the sum of the local, presynaptically-induced calcium, 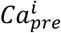, and the global, plateau-induced calcium, *Ca*_*plateau*_ .

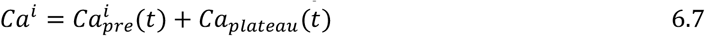

The presynaptic calcium is modeled as a pulse of calcium with an initial concentration of *pre*_*height*, which exponentially decays with a time constant of of τ_*Ca*_ :

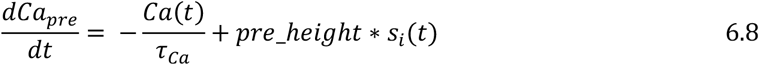

The postsynaptic calcium is modeled as a rectangular step of calcium of height *plateau*_*height* which also decays to baseline at a rate of τ_*Ca*_ once the induction ends:

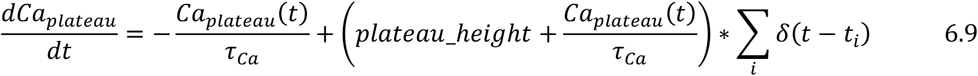

The parameters for BTSP were fitted using the differential evolution algorithm from SciPy (Virtanen et al., 2020) with a cost function designed to qualitatively reproduce the basic experimental results from (Milstein et al., 2021), (i.e. to qualitatively obtain the results shown in figure 9E-D).

## Simulation Parameters

### Frequency-dependent plasticity

Frequency-dependent plasticity:
| Parameter | Value |
| --- | --- |
| $Ca_{pre}$ | 1.05 |
| $w_0$ | 1 |
| $\tau_{Ca}^*$ | 10 ms* |
| $\theta_D$ | 1 |
| $\theta_P$ | 1.3 |
| $\eta_D$ | 0.04 |
| $\eta_P$ | 0.055 |
| $F_D$ | 0.42 (Manahan-Vaughan et al., 2000) |
| $F_P$ | 2.25 (Redondo et al., 2010) |
| $dt$ (step size for numerical integration) | 0.01 ms |

**STDP**

| Parameter | Value |
| --- | --- |
| $Ca_{pre}$ | 0.9 |
| $Ca_{post}$ | 1.55 |
| $w_0$ | 1 |
| $\tau_{Ca}$ | 7 ms* |
| $\theta_D$ | 1 |
| $\theta_P$ | 1.3 |
| $\eta_D$ | 0.1 |
| $\eta_P$ | 0.075 |
| $F_D$ | 0.42 |
| $F_P$ | 2.25 |
| $dt$ | 0.01 ms |

**BTSP**:

| Parameters |  |
| --- | --- |
| $N$ | 40 |
| $r$ | 329.23 |
| $V_{rest}$ | -75 mv |
| $V_{plateau}$ | -30 mv |
| $pre\_height$ | 0.003 |
| $plateau\_height$ | 1.295 |
| $plateau\_duration$ | 300 ms |
| $\tau_{Ca}$ | 2023.327 ms |
| $\tau_V$ | 15 ms |
| $P_{max}$ | 0.9 |
| $T$ | 10,000 ms |
| $\theta_D$ | 1.5 |
| $\theta_P$ | 2.247 |
| $\eta_D$ | 0.723 |
| $\eta_P$ | 0.002182 |
| $F_D$ | 0.06 |
| $F_P$ | 2.98 |

## Code Availability

Code for the simulations in this paper can be found at https://github.com/tmoldwin/FPLR.

## Acknowledgments

The authors would like to thank Aaron Milstein, Christine Grienberger, and Léa Caya-Bissonnette for their helpful input with respect to the BTSP portion of this paper. This work was supported by a grant from the Office of Naval Research, No. 138151-5125597, the Drahi Family Foundation, and the ETH domain for the Blue Brain Project.

---

**Supplementary Figure S1:**
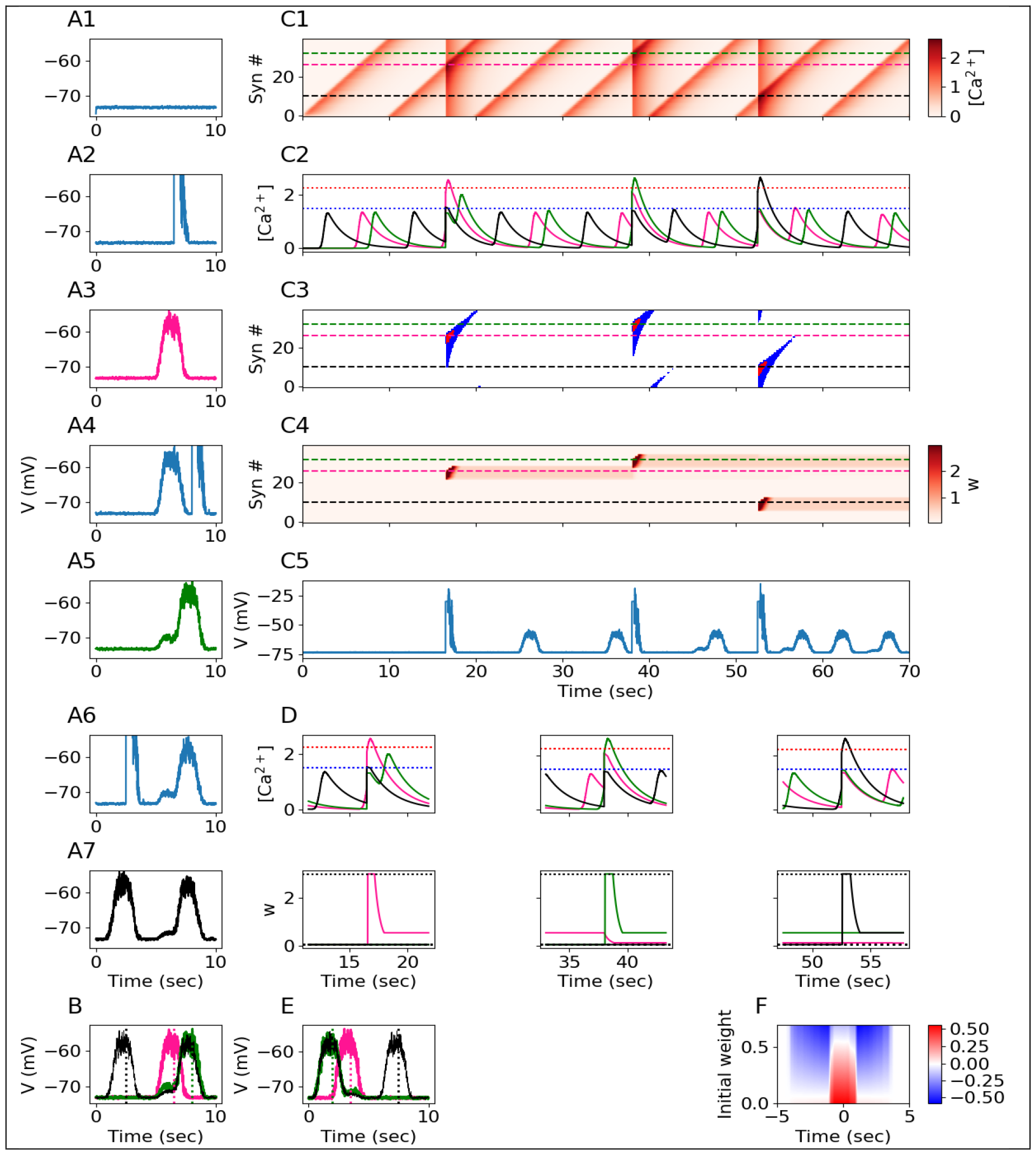
Behavioral Time Scale Plasticity “Mirror Image” Experiment (A1-A7) Voltage traces for 7 laps, each lap lasts for 10 seconds. A plateau potential is induced in the 2nd, 4th, and 6th laps. Green, pink, and black traces indicate place fields observed on the 3rd, 5th, and 7th laps, respectively. **(B)** Overlay of voltage traces for place fields from the 3^rd^ (pink), 5^th^ (green), and 7^th^ (black) laps. Vertical dashed lines indicate plateau induction location during the preceding lap. **(C1)** Total [*Ca*^2+^] per synapse (rows, 40 synapses) over the course of all 7 laps. **(C2)** [*Ca*^2+^] over time for three synapses whose receptive fields are centered at the location of the first (pink - 15) second (green) or third (black) plateau induction. Boxes show calcium traces around the time of plateau induction (see (D)). **(C3)** Plasticity bar codes for each synapse (rows) over all laps. **(C4)** Weights over time for each synapse over all laps. **(C5)** Voltage over time, as in A, for all 7 laps. **(D)** Zoom-in on the calcium traces (Top) and weights (Bottom) from the three synapses shown in B2 at the time of each plateau induction. In the second lap (left), the pink synapse’s [*Ca*^2+^] rises beyond *θ*_*P*_, inducing potentiation toward the maximum strength (dotted horizontal line at top), and the green synapse’s [*Ca*^2+^] rises above *θ*_*D*_, although the green synapse is already at the minimum strength (dotted horizontal line on bottom) so it can’t depress any further. In the fourth lap (middle), the green synapse’s [*Ca*^2+^] rises beyond *θ*_*P*_, inducing potentiation, and the pink and black synapses’ [*Ca*^2+^] rises above *θ*_*D*_, but only the pink synapse can be depressed, because it was previously potentiated. In the sixth lap (right), the black synapse’s [*Ca*^2+^] rises beyond *θ*_*P*_, inducing potentiation (the green synapse is above *θ*_*D*_ for a brief duration, but insufficiently long to substantially depress). **(E)** Result of a “mirror image” experiment, where the locations of plateau inductions are reversed (see Figure 8). Green, pink, and black traces indicate locations of inductions at the 3^rd^, 5^th^, and 7^th^ laps, as in **B. (F)** Change in weights as a function of initial weight and receptive field distance from plateau onset.

